# From spatio-temporal morphogenetic gradients to rhythmic patterning at the shoot apex

**DOI:** 10.1101/469718

**Authors:** Carlos S. Galvan-Ampudia, Guillaume Cerutti, Jonathan Legrand, Romain Azais, Géraldine Brunoud, Steven Moussu, Christian Wenzl, Jan U. Lohmann, Christophe Godin, Teva Vernoux

## Abstract

Rhythmic patterning is central to the development of eukaryotes, particularly in plant shoot post-embryonic development. The plant hormone auxin drives rhythmic patterning at the shoot apical meristem, but the spatio-temporal dynamics of the auxin gradients is unknown. We used quantitative imaging to demonstrate that auxin provides high-definition graded information not only in space but also in time. We provide evidence that developing organs are auxin-emitting centers that could self-organize spatio-temporal auxin gradients through a transport network converging on the meristem center. We further show that a memory of the exposition of cells to auxin allows to differentiate temporally sites of organ initiation, providing a remarkable example of how the dynamic redistribution of a morphogenetic regulator can be used to create rhythmicity.

## Main Text

Specification of differentiation patterns in multicellular organisms is classically thought to be regulated by gradients of morphogenetic regulators (morphogens in animals) providing positional information to cells (*1*). In plants, the hormone auxin is one of the main morphogenetic regulators (*2, 3*). This small molecule acts not only during embryonic development but also during post-embryonic development, where it is essential for the reiterative organogenesis characteristic of plants (*4*). Notably, plant shoots develop post-embryonically through rhythmic organ generation in the shoot apical meristem (SAM), a specialized tissue with a stem cell niche in its central zone (CZ). In *Arabidopsis thaliana*, as in a majority of plants, organs are initiated sequentially in the SAM peripheral zone (PZ) and at a relative angle close to 137° from the previous one, either in a clockwise or anti-clockwise spiral (*5*). SAM organ patterning or phyllotaxis has been extensively analyzed theoretically (*6-8*). A widely accepted model proposes that the time interval between organ initiations (the plastochrone) and the spatial position of organ initiation emerge from a combined action of isotropic inhibitory signals emitted by pre-existing organs and the SAM center (*6*). Tissue growth would then self-organize organ patterning by moving organ-associated signaling centers away from the stem cells and leaving space for new ones.

Biologically, evidence suggests that auxin provides the positional information that drives phyllotaxis patterning (*9, 10*). Auxin, thought to be synthesized throughout the meristem (*11-14*), has been proposed to be transported directionally toward incipient primordia where it activates a transcriptional response leading to organ specification (*3, 9, 15*). A network of PIN-FORMED 1 (PIN1) efflux carriers, whose polarity determines the direction of auxin fluxes, regulates auxin spatio-temporal distribution cooperatively with other carriers (*9,16*). The activity of this network results in accumulation of auxin that triggers organ initiation. The PIN1 network was also proposed to create an auxin depletion at the organ periphery that specifies organ boundaries and blocks organ initiation in the organ vicinity (*9,17-21*). In addition, it has been shown that the CZ is markedly less responsive to auxin (*17, 18*). Altogether, these regional cues restrict new organ location in the growing SAM as proposed in formal models. The genetically-encoded biosensor DII-VENUS, a synthetic protein degraded directly upon sensing of auxin, recently provided a first qualitative visualization of spatial auxin gradients in the SAM (*17, 22*).

The SAM is rather unique in that it implicates a continuous redistribution of a morphogenetic regulator in a growing tissue with helicoidal symmetry. This suggests that auxin could carry spatio-temporal morphogenetic information in the SAM. This is reminiscent of recent findings in animals that are questioning whether morphogenetic signals carry information only in space (as originally proposed in the morphogen concept (*23*) and suggests rather a spatio-temporal nature for positional information (*24-26*). Here, we used a quantitative imaging approach to reveal that auxin indeed provides spatio-temporal morphogenetic information, analyze the mechanisms generating auxin 4D dynamics and understand how this information is processed in the SAM to generate rhythmic patterning.

## Results

### Spatio-temporal auxin distribution

In the SAM, the biosensor DII-VENUS fluorescence reports for auxin concentration with cellular resolution (*17, 22*). To extract quantitative data on auxin distribution, we generated a DII-VENUS ratiometric variant, named hereafter qDII (quantitative DII-VENUS). qDII consists of a RPS5A promoter driving stoichiometric co-expression of DII-VENUS and a non-degradable TagBFP reference (*27, 28*)(Fig. S1A-H). We also introduced in plants expressing qDII a stem cell-specific CLV3::mCherry nuclear transcriptional reporter (*29*) that provided a functional and robust geometrical reference of the SAM center (Fig. 1A-B and Fig. S1I-M).

**Figure 1.**
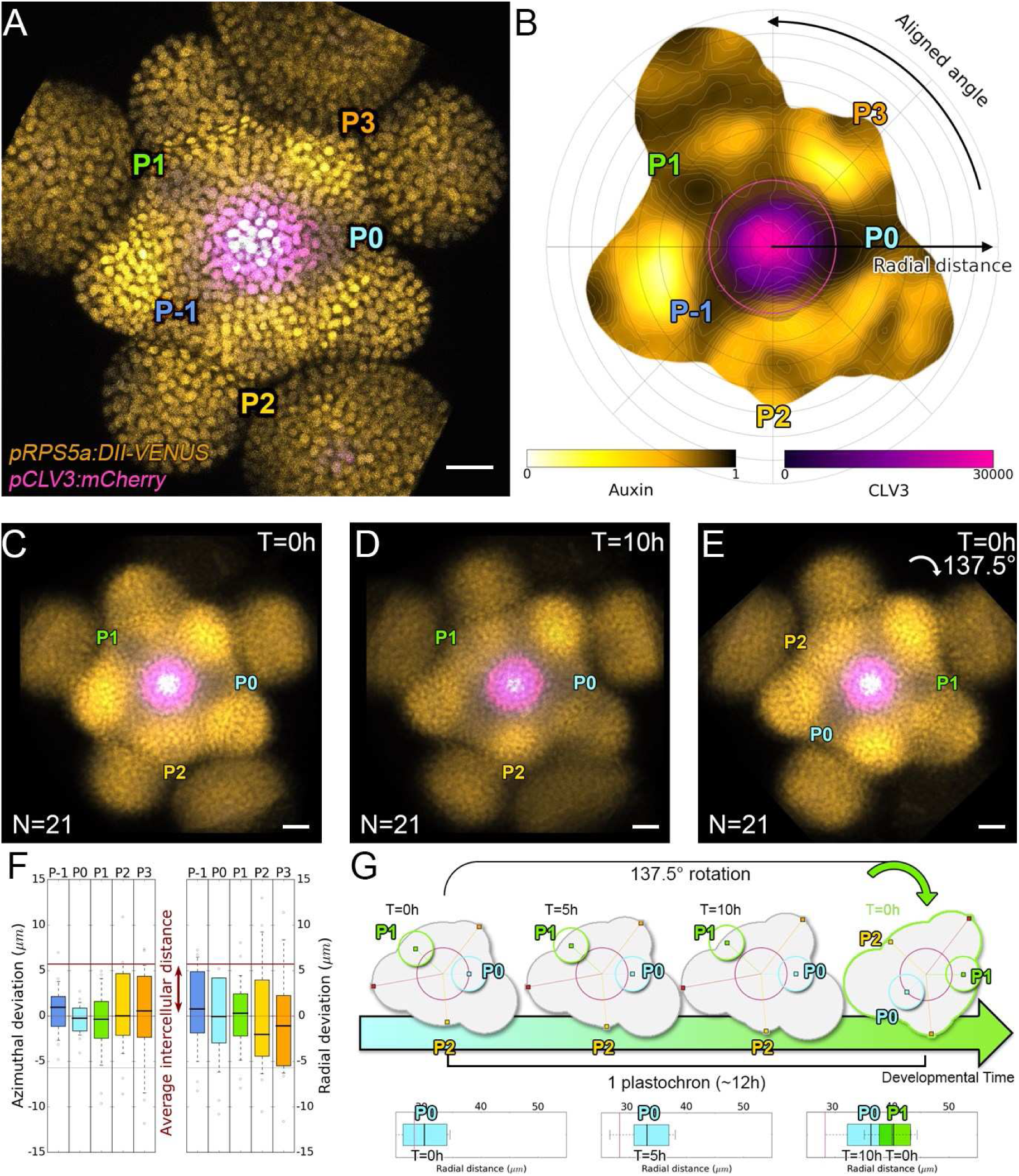
Spatial auxin distribution in the SAM follows a precise and reiterative pattern. **A.** Representative qDII expression pattern (VENUS-N7 in yellow and nuclei in grey) and CLV3 transcriptional reporter line (magenta). Primordia are indicated by colour and rank **B.** Auxin map (yellow) of (**A**) showing CLV3 radial extension (circle in magenta). Black arrows depict two main parameters for registration (radial distance from the centre and aligned angle). **C-E.** Superposition of 21 registered SAM images at time 0 h (**c**), 10 h later **(D).** E. 137.5° clockwise rotation of **(C)** results in a quasi-identical image of **(D).** See **Figure S2A** for non-registered image superposition. Scale bars = 20 μm. **F.** Precision in auxin maxima positioning. Distance to a reference position in both azimuthal direction (angle in a Fibonacci phyllotactic spiral, left panel) and radial direction (population average trend, right panel). Red lines indicate the average cellular distance. N = 21 meristems. Colours indicate the auxin maxima primordium rank (P_-1_ blue, P_0_ cyan, P_1_ green, **P**_2_ yellow, **P**_3_ orange) **G.** Space can be used as a proxy for time, as a rotation of 1 divergence angle is an equivalent to a translation of 1 plastochrone (12h) in time.

All analyzed meristems (21 individual SAM, 24945 nuclei) showed qDII patterns similar to the one obtained with DII-VENUS, with auxin maxima locations following the phyllotactic pattern (*17*) (Fig. 1A-B). Despite the fact that SAMs were imaged independently and not synchronized, qDII patterns appeared highly stereotypical with easily identifiable fluorescence maxima and minima. This was confirmed by image registration using SAM rotations (applying prior mirror symmetry if necessary; Fig. 1C-D and Fig. S2A-C). All images could be superimposed with limited loss of information definition (Fig. S2A-C). This shows that auxin distribution follows the same synchronous pattern at population scale, with restricted angular and rhythmicity variability (Fig. S2F-G, Supplementary Method 1), resulting, up to a rotation, in apparent stationarity (Fig. 1G).

To further quantify auxin distribution, we developed a mostly automated computational pipeline to extract SAM quantitative fluorescence (Fig. 1B and Supplementary Method 2). We used the spatial distribution of 1-DII:VENUS/TagBFP as a proxy for auxin distribution, named hereafter “auxin” (Fig. 1B) and focused on the epidermal cell layer (L1) where organ initiation takes place (*30*). Primordium 0 (P_0_) was defined as the location of the absolute auxin maximum in the PZ and other local maxima where called P_n_ (Supplementary Note 1), with n corresponding to their rank in the phyllotactic spiral. The pipeline then allows quantifying nuclear signal information and aligns all the SAMs to a common clockwise reference frame with standardized x,y,z-orientation with the P_0_ maximum to the right. This automatic registration showed that auxin maxima are positioned in the SAM with a precision close to the size of a cell both in distance to the center and in azimuth (with a maximal standard deviation of 8.4 μm or 1.5 cells, Fig. 1F).

We then considered the temporality of auxin distribution by using time-lapses over a time span close to the system period, the plastochrone. P_0_ and successive auxin maxima moved radially (Fig. S2D). Remarkably, while the average radial distance of each local maximum P_n_ to the SAM center progresses (Fig. S2D), their standard deviation does not change significantly in time, reflecting the synchronized movement of local maxima, with limited meristem to meristem variation. After 10h, every P_n_ local maximum has almost reached the starting position of the next local maximum, P_n+1_, but after 14h they have passed it over (Fig. S2D). This suggests that a rotation of 137.5° (Fig. S2E) that replaces P_n_ by P_n+1_ corresponds to a 10 to 14 hours temporal progress (Fig. 1G). This is confirmed by error measures obtained with different rotation angles (Fig. S2H), allowing to conclude that plastochrones have a value of 12h ± 2h. We thus derived a continuum of primordium development by placing P_n+1_ time series one plastochrone (12h) after P_n_ time series on a common developmental time axis (Fig. 1g). Together with the developmental stationarity, this allows reconstructing auxin dynamics over several plastochrones from observations spanning only one. The quantitative temporal map of auxin distribution in the SAM obtained through this reconstruction demonstrates dynamic building of auxin maxima in the PZ first as finger-like protrusions (visible at P_-2_, P_-1_ and P_0_) from a permanent high auxin zone at the center of the meristem (Movie S1), as previously predicted (*18*). At later stages, auxin minima are progressively established precisely in between the auxin maxima and the CZ and not surrounding the auxin maximum (Fig. S3).

We next wondered whether the motion of auxin maxima and minima could purely result from cellular growth as proposed in previous theoretical models (*6, 19, 31*). Following a P_1_ maximum, we observed that cells in the auxin maximum at time 0h gradually became part of the depletion zone at time 10h (Fig. 2A-C). Using nuclear motion to estimate cell motion vectors and compare them with auxin maxima motion, we further found that the average radial speed of auxin maxima between stages P_1_ and P_4_ surpasses the average displacement of individual nuclei, by up to more than 1μm/h (or nearly 2 cells in 10h) at its peak in P_2_ stage (Fig. 2D-E). These results show that auxin maxima are not attached to specific cells; instead they travel as a wave in the tissue. Consequently, the SAM cellular network provides a dynamic medium in which auxin distributions move radially with their own velocity relative to the growing tissue (Fig. 2D-E). Analysis on time-courses up to 14h revealed significant auxin variations in certain cells over one plastochrone while auxin levels remained unchanged in others (Fig. 2F and 2G, bottom rows). However, neighboring cells always showed limited differences in their temporal auxin profiles. Altogether, we concluded from these observations that there is a high definition spatio-temporal distribution of auxin that moves faster than growth in the tissue and provides cells with graded morphogenetic information both in space and time (Fig. 2H).

**Figure 2.**
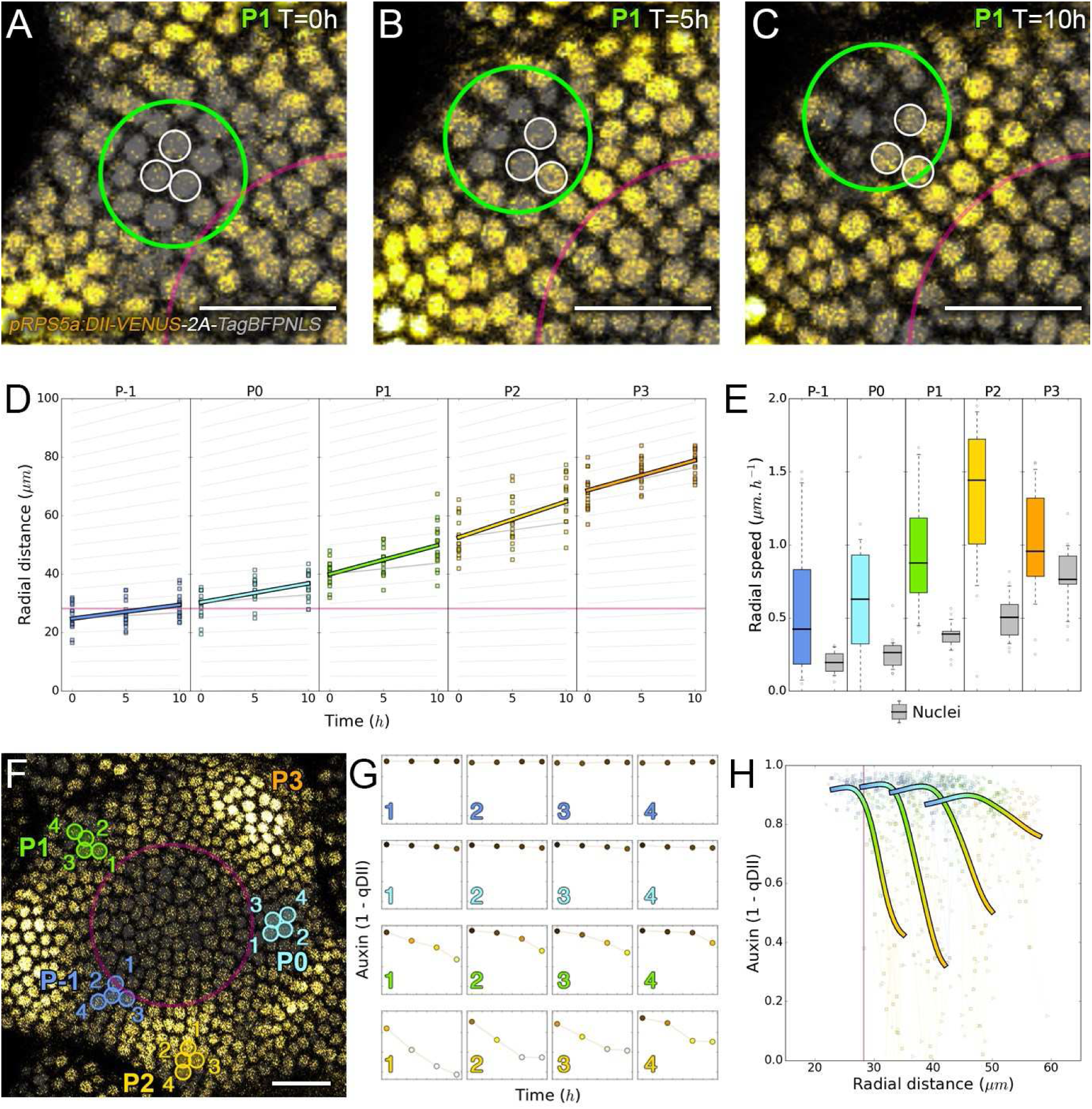
Auxin dynamics are not cellularly fixed. **A-C.** Representative projection of P_1_ nuclei in time showing qDII (grey and yellow). Time tracked nuclei are marked by a white circle showing rapid changes of auxin in 10 h. Green circle represents the position of the auxin maxima at each time point. The magenta line indicates the CZ. Scale bars = 20 μm. **D.** Average motion of maxima (colour lines) is faster than average cell motion (grey lines). The magenta line indicates the CZ border. N = 21 meristems. **E.** Compared distributions of radial motion speeds of auxin maxima estimated as the slope of a linear regression per individual and individual nuclei radial speed at the location of the maxima. N = 21 meristems. **F.** Individual cells experience different auxin histories. Tracked cells at different locations (coloured circles). Scale bar = 20 Δm. **G.** Corresponding auxin levels (ordinate) in time (abscissa 0, 5, 10, 14 h) of tracked cells in **(F). H.** Cellular mean auxin trajectories as a function of radial distance. Each line represents an extrapolated cell-size sector moving accordingly to cellular radial motion by its Gaussian average trajectory in radial distance (abscissa) and auxin value (ordinate). The colour indicates the developmental stages at a given radial distance (P_-1_ = blue, P_0_ = cyan, P_1_ = green, P_2_ = yellow).

### The control of auxin spatio-temporal dynamics

The creation of auxin maxima first as protrusions of a high auxin zone in the CZ contrasts with the largely prevailing vision of organogenesis being triggered by local auxin accumulation at the periphery of the CZ and concomitant auxin depletion around auxin maxima (*9, 18-21*). This, in addition to the partial uncoupling of auxin distribution dynamics and growth, led us to question our current knowledge of the spatio-temporal patterns of PIN1, given their central role in controlling auxin distribution (*9, 18, 19*, 21). Co-visualization of a functional PIN1-GFP (*3*) and qDII/CLV3 fluorescence over time showed that PIN1 concentration increases from Po and reaches a maximum at P_2_ before decreasing (Fig. 3A-C), consistently with previous observations (*15, 32, 33*). To quantify PIN1 cellular polarities, we used staining with the fluorescent dye propidium iodide (PI) as a reference to position the PIN1-GFP signal in 3D relative to the L1 anticlinal cell walls (*34*) (Fig. 3B and Supplementary Method 2). These 3D high-resolution reconstruction of PIN 1 polarities demonstrated that the crescent-shape often thought to indicate polarities in cells does not always correlate with polarities and can thus be sometimes misleading (Fig. S4). We mapped the vector fields of polarities to identify the trends in auxin flux directions at the scale of the SAM (Fig. 3D, and Supplementary Method 2). While some local converging PIN1 polarities can be seen close to auxin maxima at the PZ (Fig. 3E-F), the vector fields rather show large-scale convergence of PIN 1 toward the center of the SAM that meet in front lines where auxin maxima protrude from the CZ (Fig. 3D and Fig. S5). We detected the inversion of PIN1 polarities at organ boundaries previously observed (*15*) and our quantifications show that this occurs only from P_6_, thus isolating the flower from the rest of the meristem from this late stage. P_3_ to P_5_ stages show a general flux toward the SAM that is locally deflected around the zones of auxin minima before converging back toward the meristem center (Fig. 3D and Fig. S5). Over the course of a plastochrone, only limited changes of the PIN1 network are observed (Fig. S5), suggesting that changes in auxin distribution at this time resolution might not require important adjustments of the direction of the auxin fluxes. Other carriers are active in the SAM (*16*) and the role of PIN 1 in auxin distribution could also have been overestimated.

**Figure 3.**
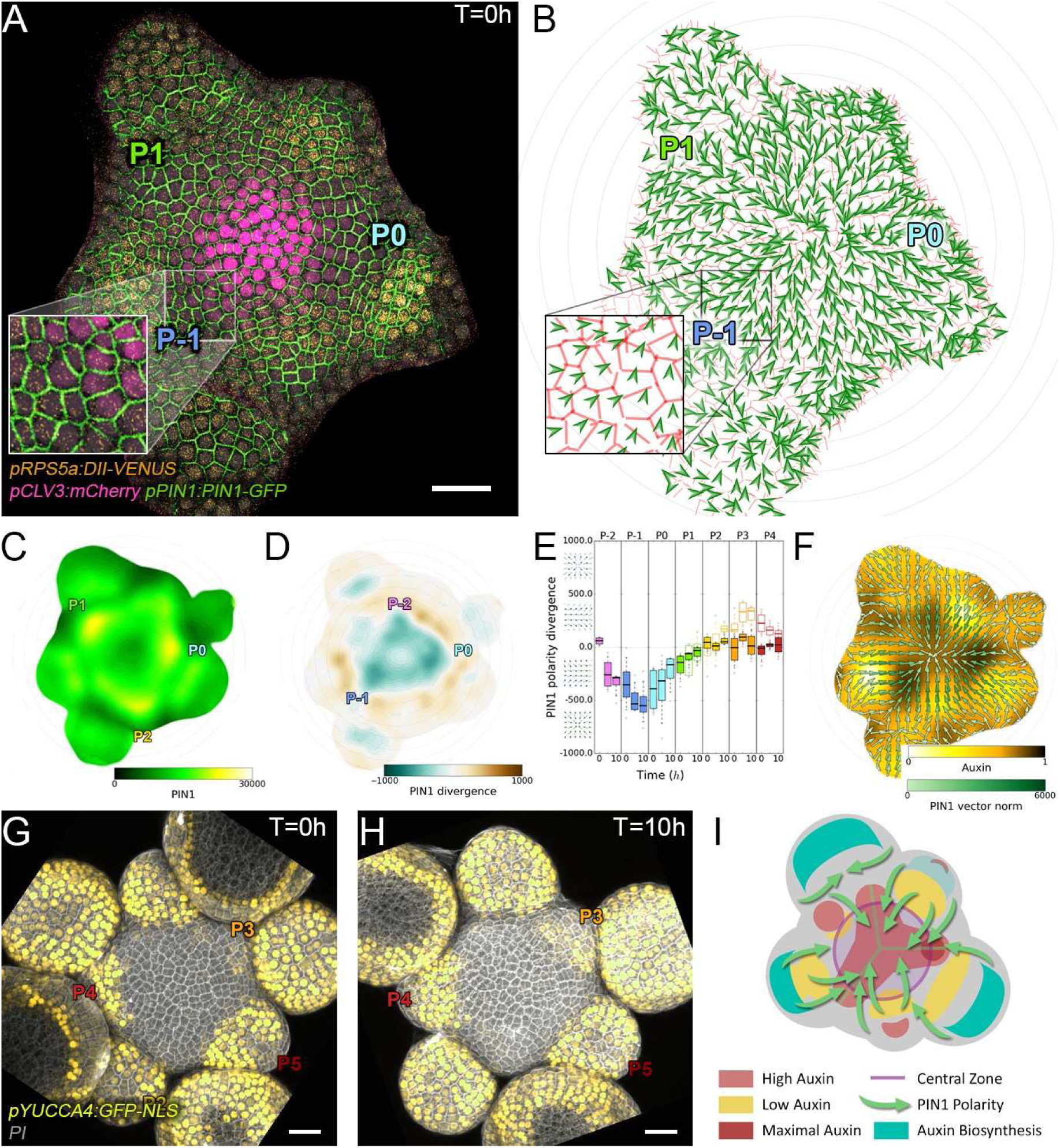
Auxin polar transport and biosynthesis act synchronously to regulate patterning. **A.** PIN1 expression patterns follow auxin dynamics. Co-visualization of qDII (yellow), CLV3 (magenta) and PIN1-GFP (green) in time. Scale bar = 20 μm. Inset shows a magnification of P_-1_ sector **B.** Cellular main polarity vectors of (**A)** detected by our method (see supplementary method 2). Inset shows a magnification of P_-1_ sector. **C.** PIN1-GFP expression map shows accumulation at P_0_, P_1_ and P_2_ auxin maxima. **D.** PIN1 convergence points are located at P_-1_ and P_0_. Polarity divergence map indicating convergence (blue), parallel alignment (white) or divergence (orange). **E.** Convergence of PIN1 is transient and localized to P_-1_/P_0_. PIN1 polarity local divergence values at the auxin maxima (colour filled boxplots) or auxin minima (white filled boxplots). Boxplot centres show median. **F.** PIN1 is mainly polarized towards the CZ. Vector fields were generated from PIN1 cell polarities (green arrows). Convergence of vectors occurs at auxin maxima. **G-H.** Activation of auxin biosynthesis at P_3_/P_6_ feeds the SAM of auxin. YUC4:GFP transcriptional reporter is activated at the epidermis cell layer of primordia. Scale bars = 20 μ m. **I.** Transport front lines and biosynthesis tissue memories control the formation of auxin protrusions.

The analysis of the transport network led us to question where auxin could be produced in the SAM. YUCCAs (YUCs) have been shown to be limiting enzymes for auxin biosynthesis during development (*11, 35*). We thus mapped expression of the eleven YUCs in the SAM (Fig. S6)(*35, 36*). Only *YUC1,4,6* were detected and only *YUC6* in the SAM proper with a very weak expression in the CZ (Fig. 3G-H and Fig. S6). This is coherent with genetic and expression data (Table S1) (*11*) Both *YUC1* and *YUC4* are expressed at the L1 layer on the lateral sides of the SAM/flower boundary from P_3_ for *YUC4* and in P_4_ for *YUC1* (*11*) (Fig. 3G-H and Fig.. S6). From P_4_, *YUC4* expression extends over the entire epidermis of flower primordium. Together with the PIN 1 network organization, these expression patterns thus identify the primordia at P_3_-P_5_ stages as auxin production centers for the SAM.

In conclusion, there is a structured global organization of the PIN1 pump network, with high concentrations of auxin at the center of the SAM but also at P_-1_ and P_0_, acting as flux attractors and with auxin originating from developing flowers at P3-P5 stages (Fig. 31).

### The role of time in transcriptional responses to auxin

To assess quantitatively whether and how auxin spatio-temporal distribution is interpreted in the SAM, we next introduced the synthetic auxin-induced transcriptional reporter DR5 (*37–39*) driving mTurquoise into the qDII/CLV3 reporter line (Fig. 4A-D). Cells expressing DR5 closest to the CZ were robustly positioned at an average distance of 32 μM ± 7 (SD) from the center. This corresponds to a distance at which the CLV3 reporter expression is lower than 5% of its maximal value (Fig. S7A). The distance from the center at which transcription can be activated by auxin is thus defined with a near-cellular precision.

**Figure 4.**
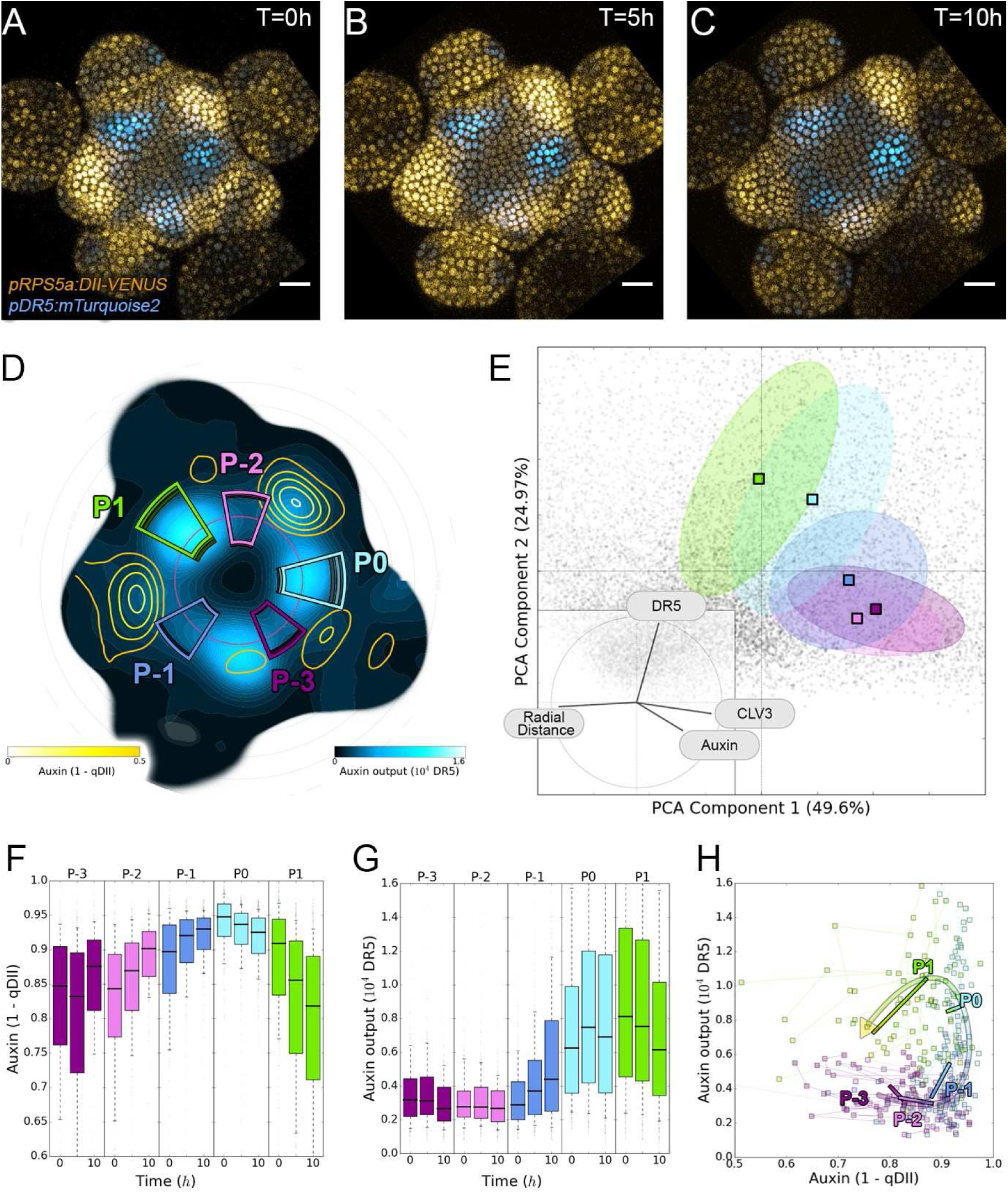
Auxin and its transcriptional output show a complex non-linear relationship. **A-C.** Time lapse images of representative projections of qDII (yellow) and DR5 auxin-activated synthetic promoter (cyan). Scale bars = 20μ m. **D.** DR5 expression map (cyan) showing auxin depletion zones (yellow contour lines). Coloured sectors show the tissue areas where primordia are located (P_-3_ to P_1_) moving according to cellular motion with a linear radial deformation of the tissue. **E.** Principal Component Analysis (PCA) showing absence of correlation between auxin and DR5 at global scale. Coloured ellipses show the consistent pattern associated with each primordium stage. **F-G.** DR5 and auxin waves are delayed by nearly a plastochron. Coloured boxes correspond to the regions shown in **D.** Boxplot centres show median. **H.** Auxin and DR5 non-linear relationship. Cells at different spatial loci (indicated by colours) can have the same auxin input but trigger different transcriptional responses. Lines represent the regression of auxin and DR5 medians in time.

To obtain a global vision of how auxin transcription is related to auxin concentration, we performed a Principal Component Analysis (PCA) using quantified levels of DR5, auxin and CLV3 in each nucleus of the PZ of 10h time series together with their distance from the center (Fig. 4E). With the first two axis reporting for around 75% of the observed variability, we unexpectedly observed orthogonality between auxin input and DR5 output clearly marking an absence of a general correlation in the meristem (Fig. 4E, inset). This unexpected finding was confirmed by the low numerical values of Pearson correlation coefficients between DR5 and auxin values at cell-level (Fig. S7B). We refined our analysis by studying locally correlations between auxin and DR5 in the different primordia regions (Fig. 4D). We observed homogeneous auxin and DR5 behaviors that characterized each region (Table S2), showing that the spatial position of a cell conditions the link between its auxin input and the state of its transcriptional response. We then assembled on a single graph all the observed couples of values (auxin, DR5) averaged over each primordium region (Fig. 4H). This evidenced that, spatially, a given auxin value does not in general determine a specific DR5 value. However, values corresponding to primordia at consecutive stages follow loop-like counter-clockwise trajectories in the auxin x DR5 space (indicated by an arrow in Fig. 4H). Such trajectories are symptomatic of hysteresis reflecting the dependence of a system on its history. In other words, DR5 expression is dependent on the developmental history of the cells.

We then used our reconstructed continuum of primordium development to study the joint temporal variations of DR5 and auxin within a local group of cells during initiation (Supplementary Method 3). This showed that the start of auxin-induced transcription follows the building-up of auxin concentration with a delay of nearly one plastochrone (Fig. 4F-G). The duration of the observed phenomenon suggests the existence of an additional process, different from protein maturation (*40*), that creates a significant DR5 response delay of primordia cells to auxin during development. Due to this delay, DR5 is not a direct readout of auxin concentration, explaining in these cells the absence of correlation between DR5 expression and auxin levels.

We next wondered what could explain a time-dependent acquisition of cell competence to respond to auxin. A first possible scenario is that cells exiting the CZ proceed through different stages of activation of an auxin-independent developmental program enabling them to sense auxin after a temporal delay. A second possibility is that auxin controls this developmental program through a time integration process. In that case, cells exiting the CZ would need to be exposed to high auxin concentration for a certain amount of time to mount up an auxin transcriptional response. To test these scenarios, we treated SAMs with auxin for different durations (Fig. 5A-I). All treatments equally saturated auxin levels in the SAM (Fig. S8A). In the shorter auxin treatments (30’ and 120’), auxin output was only enhanced at P_0_, P_1_ and P_2_ i.e. where cells have already been exposed to auxin (Fig. 5F-G and I, Fig. S8B). On the other hand, the longer auxin treatments (300’) activated signaling in most cells in the PZ and organs (Fig. 5H-I). Both observations are compatible with the second scenario where longer exposure allows activating signaling in more cells and are incompatible with the first one, where the capacity of the cells to respond to auxin is intrinsic and does not dependent upon auxin exposure time. Our results thus indicate that temporal integration of the auxin signal controls the activation of transcription in the SAM.

**Figure 5.**
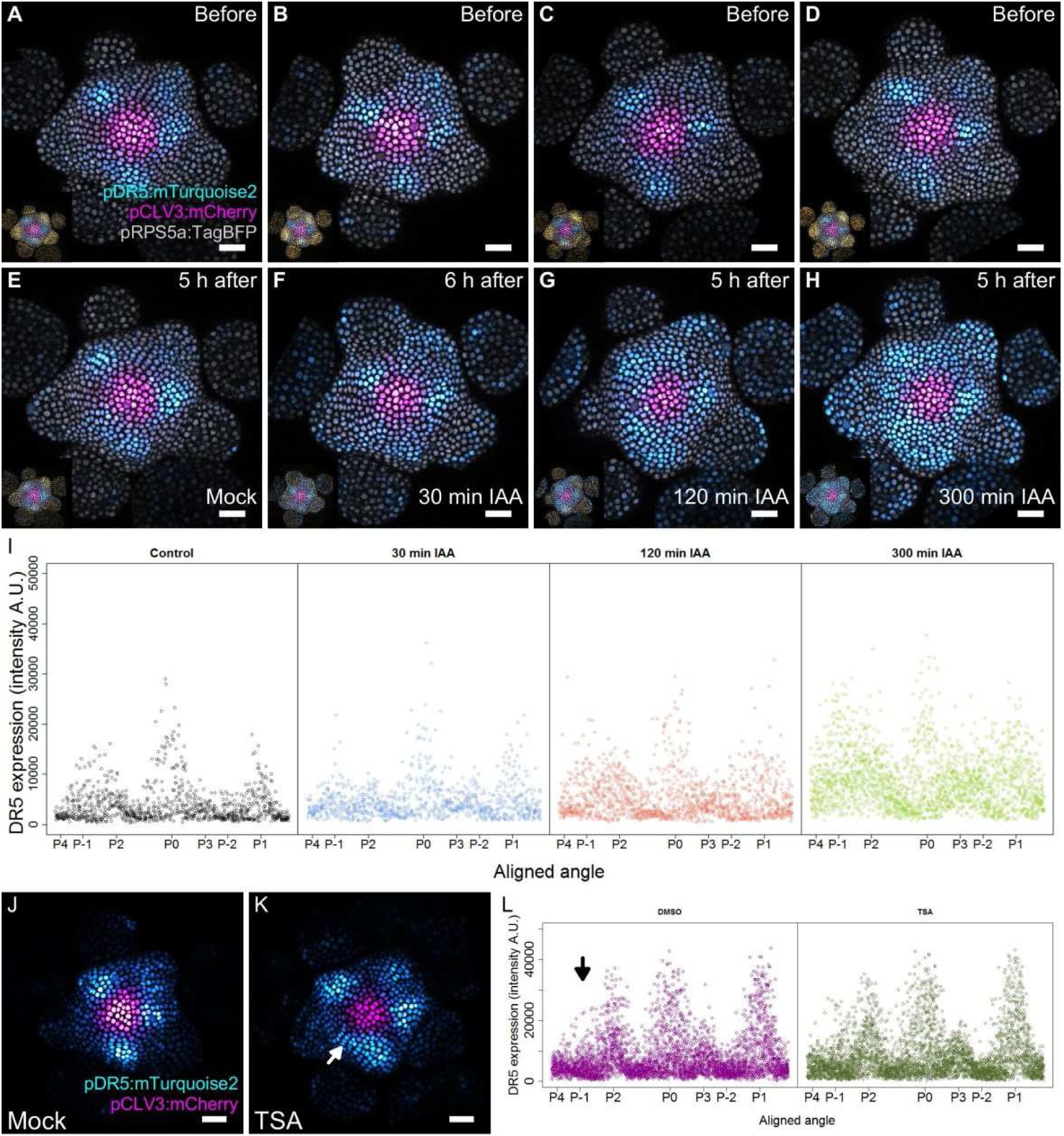
Signal temporal integration regulates auxin output through epigenetic control. **A-D.** Representative images of DR5 expression before auxin treatment. **E.** Mock treated meristem after 5 h. **F-H.** Images of meristems after 5 or 6 h treatment. Meristems were treated with 1 mM IAA for 30 min **(F),** 120 min **(G)** or 300 min **(H).** DR5 (cyan), pRPS5a (gray) and CLV3 (magenta) labelled nuclei are shown. qDII (yellow) of the same image is shown in the inset, **i.** Quantification of DR5 expression at the PZ after auxin treatments. Each dot represents a nuclei (Mock N=3045, 30 min N=1895, 120 min N=3378, 300 min N= 5081). The angular location of primordia are indicated on the abscissa. **J-k.** Representative image of DR5 of a meristem treated with mock or the histone deacetylase inhibitor TSA. White arrow indicates the region P_-1_. **L.** Quantification of DR5 expression in the PZ epidermal cell layer of **J** and **K.** Each dot represents a nuclei (Mock N=5621, TSA=5585). The angular location of primordia is indicated. Black arrow indicates the region P_-1_. Scale bars = 20μ m.

Chromatin state is one mechanism that allows for temporal integration of signals (*24, 41–43*). Also, auxin signaling have been shown to act by modifying acetylation of histones (*44, 45*). Pharmacological inhibition of histone deacetylases (HDACs) alone was able to trigger concomitant activation of DR5 at P_0_ and P_-1_ sites in the PZ (Fig. 5J-L, Fig. S8C). This result suggests that the timing of auxin signaling induction at the P_-1_ site depends on the chromatin acetylation status, providing a molecular mechanism for temporal integration of auxin-based positional information.

## Discussion

In a recent modeling work, the analysis of mutants perturbed in their plastochrone led to postulate a stochastic mechanism for organ initiation based on temporal integration of morphogenetic signals (*46*). Here we provide evidence that organ initiation in the SAM is indeed dependent on temporal integration of the auxin signal. Our quantitative analysis of the dynamics of auxin distribution and response supports a scenario where rhythmic organ initiation at the SAM is driven by a combination of high-precision spatio-temporal graded distributions of auxin and of the use of the duration of cell exposition to auxin to differentiate temporally sites of organ initiation (Fig. 6). Such a mechanism is likely essential for rhythmic organ patterning in the SAM as auxin-based spatial information pre-specifies several sites of organ initiation and is thus likely insufficient (Movie S1).

**Figure 6.**
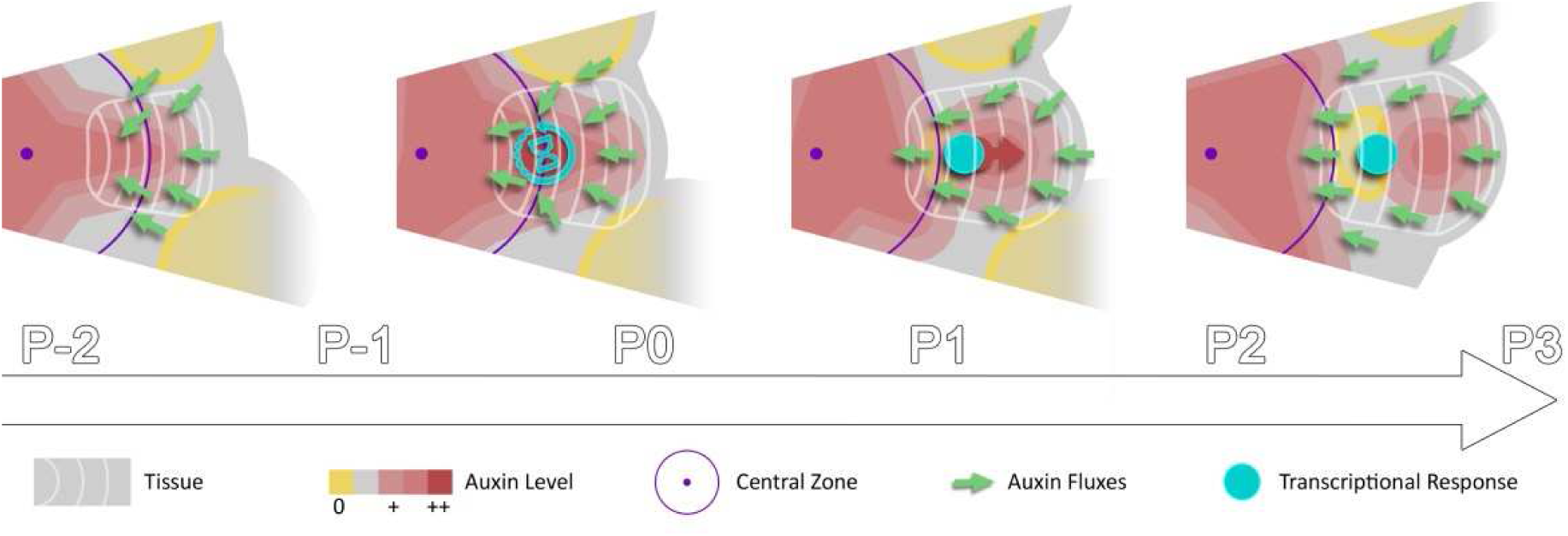
Spatio-temporal gradients of auxin translate into rhythmic organ patterning through time integration. A maximum of auxin protrudes from a high auxin concentration zone at the CZ faster than the cells move radially, possibly as a result of centripetal auxin fluxes. Cells exiting the CZ that are exposed to high auxin levels progressively acquire competence for transcriptional response through chromatin-mediated time-integration of the auxin signal. This leads to activation of transcriptional responses with a delay close to the system period, the plastochrone.

Temporal integration of the auxin signal could occur at the boundary of the CZ through a chromatin acetylation mechanism, the chromatin acetylation status of cells acting as a memory of their exposition time to auxin. Activation of transcription downstream of auxin by the Auxin Response Factors (ARF) has recently been shown to rely on chromatin remodeling, increasing the accessibility of ARF targets and possibly allowing for the recruitment of histone acetyltransferases (*45*), together with the release of histone deacetylases (HDACs) from target loci by degradation Aux/IAA repressors. Indeed, Aux/IAAs have the capacity to bind ARFs and to recruit indirectly HDACs that are thought to mediate the repression of the target loci in the absence of auxin (*44*). It is thus likely that time integration of the auxin signal in cells leaving the CZ is set by progressive acetylation of histones triggered by ARFs on their target loci. As chromatin deacetylation represses auxin signaling in the CZ (*47*), balancing the acetylation status of ARF target loci provides an original mechanism tightly linking stem cell maintenance and differentiation by precisely positioning organ initiation at the boundary of the stem cell niche, while at the same time allowing for sequential organ initiation.

The existence of high definition spatio-temporal auxin gradients suggests that similarly to several morphogens in animals (*24, 48–50*) the robustness of SAM patterning is at least in part due to highly reproducible spatio-temporal signal distribution. Our results indicate that auxin maxima could first emerge from the CZ at the confluence of centripetal auxin fluxes. Confluences creating auxin maxima would at the same time diverge fluxes away from areas where auxin minima appear (Fig. 31). Our analysis questions how auxin transport could generate this high definition signal distribution and whether different models that have been proposed can explain this distribution (*16, 20, 21, 51, 52*). Further analysis of the control of auxin spatio-temporal distribution needs also to consider that early flowers at specific stages could be auxin production centers. These flowers not only provide a memory of the developmental pattern through lateral inhibition but also through contributing positively to a self-sustained distribution of auxin by redistributing auxin back in the system (Fig. 3I). Also, our work suggests that the stem cell niche could play the role of a system-wide organizer of the auxin transport network, coherently with previous work (*18*). This could provide another layer of regulation tightly coordinating differentiation to the presence of a stem cell niche mostly insensitive to auxin (*17, 47*). The analysis framework we have established and the extensive quantitative resource we have generated will certainly be crucial to further understand how robust spatio-temporal distribution of auxin are achieved in the meristem.

## Acknowledgements

We thank Fabrice Besnard and the members of the SIGNAL team for insightful discussions; Antoine Larrieu for helping with RNA-Seq analysis; Hélène Robert-Boisivon for the YUC transcriptional lines; Olivier Hamant, Arezki Boudaoud and Jan Traas for feedback on the manuscript. We acknowledge the contribution of SFR Biosciences (UMS3444/CNRS, US8/Inserm, ENS de Lyon, UCBL) facilities PLATIM, for assistance with microscopy. This work was supported by Human Frontier Science Program organization (HFSP) Grant RPG0054-2013 to T.V. and C.G., ANR-12-BSV6-0005 grant (AuxiFlo) to T.V. and DFG FOR2581 to J.L.

## Author Contribution

C.G. and T.V. designed the project; C.G.-A., G.C., J.U.L., C.G. and T.V. designed experiments; C.G.-A., G.C., J.L., R.A., G.B., S.M., C.W. performed experiments; C.G.-A., G.C., J.L., R.A., G.B., S.M., C.G. and T.V. were involved in data analysis; C.G.-A., G.C., C.G. and T.V. wrote the manuscript with inputs from all authors.

## Data and software availability

All experimental data and quantified data that support the findings of this study are available from the corresponding authors upon request.

Generic quantitative image and geometry analysis algorithms are provided in Python libraries timagetk, cellcomplex and tissue_nukem_3d (https://gitlab.inria.fr/mosaic/) made publicly available under the CECILL-C license. Specific SAM sequence alignment and visualization algorithms are provided in a separate project (https://gitlab.inria.fr/gcerutti/sam_spaghetti.git). All other custom source codes and analysis scripts are available from the corresponding authors upon request.

## Supplementary Materials

Materials and Methods

Figures S1-S8

Tables S1-S2

Caption for Movie S1

Supplementary Notes 1. Definition of a conceptual frame for models and analysis

Supplementary Methods 1. Effects of variability on a theoretical phyllotactic system Figure S9

Supplementary Methods 2. Computational pipelines for image and data analysis Figures S10-12

Supplementary Methods 3. Extrapolated cell motion in the developmental continuum Figure S13

References (1-19)

### Materials and Methods

Seeds were directly sown in soil, vernalized at 4 °C, and growth for 24 days at 21 °C under long day photoperiod (16 hrs light, LED 150μmol/m^2^/s). Shoot apical meristems from inflorescence stems with a length between 0.5 and 1.5 cm where dissected and cultured *in vitro* as described in (***1***) for 16 hrs. When required, meristems were stained with 100 μM propidium iodide (Sigma) for 5 min. Auxin treatments were performed by immersing meristems with 1 mM Indole-acetic acid (IAA) for different times. Trichostatin A (TSA - Invivogen) was added to the ACM plates to a final concentration of 5 μM. Meristems were cultured in TSA for 16 hrs prior auxin treatment. For time lapses, first image acquisition (T=0) correspond to 2 hrs after the dark period, with the exception of auxin treatments where imaging is right after lights were on.

Previously published transgenic lines used in this study are PIN1-GFP (***2***), promCLV3:mCherry-NLS **(*5*),** and promYUC1 to 11-GFP ***(4, 5*).** Quantitative DII (qDII), promRPS5a:DII-VENUS-N7-p2A-TagBFP-SV40, reporter line and the auxin synthetic promoter DR5rev:2x-mTurqouise2-SV40 was cloned using Gateway technology (Life Sciences), and transformed in *Arabidopsis thaliana* (Col-0). Stable qDII homozygous lines were then crossed with promCLV3, promDR5rev:2x-mTurqouise2-SV40 and PIN1-GFP reporter lines.

### Microscopy

All confocal laser scanning microscopy was done with a Zeiss LSM 710 spectral microscope. Multitrack sequential acquisition was performed using always the same settings (PMT voltage, laser power and detection wavelengths) as follows: VENUS, excitation wavelength (ex): 514 nm, emission wavelength (em): 520-558 nm; mTurquoise2, ex: 458 nm, em: 470-510 nm; EGFP, ex: 488 nm, em: 510-558 nm; TagBFP, ex:405 nm, em: 430-460 nm; mCherry, ex: 561 nm, em: 580-640 nm; propidium iodide, ex: 488, em: 605-663 nm.

### Quantification and statistical analysis

All confocal images were pre-processed using the ImageJ software (http://rsbweb.nih.gov/ij/) for the delimitation of the region of interest. Then the CZI image files were processed in a computational pipeline developed by the authors and relying essentially on the numpy, scipy, pandas, czi_file Python libraries, as well as other custom libraries. Extensive details about the developed methods and algorithms are given in Supplementary Method 2.

Confidence intervals were calculated with a confidence level of 95% in the R environment (***6***). The boxplots displayed in the article were obtained by computing the median (central line), first and third quartiles (lower and upper bound of the box) and first and ninth deciles (lower and upper whiskers) using the R environment or numpy percentile function and rendered using the matplotlib Python library. Linear regressions were performed using the polyfit and polyval numpy functions. P-values were obtained using the scipy anova implementation in the f_oneway function. Principal component analysis was performed using the PCA implementation from the scikit-learn Python library. All data were generated with at least 3 independent sets of plants.

**Figure S1.**
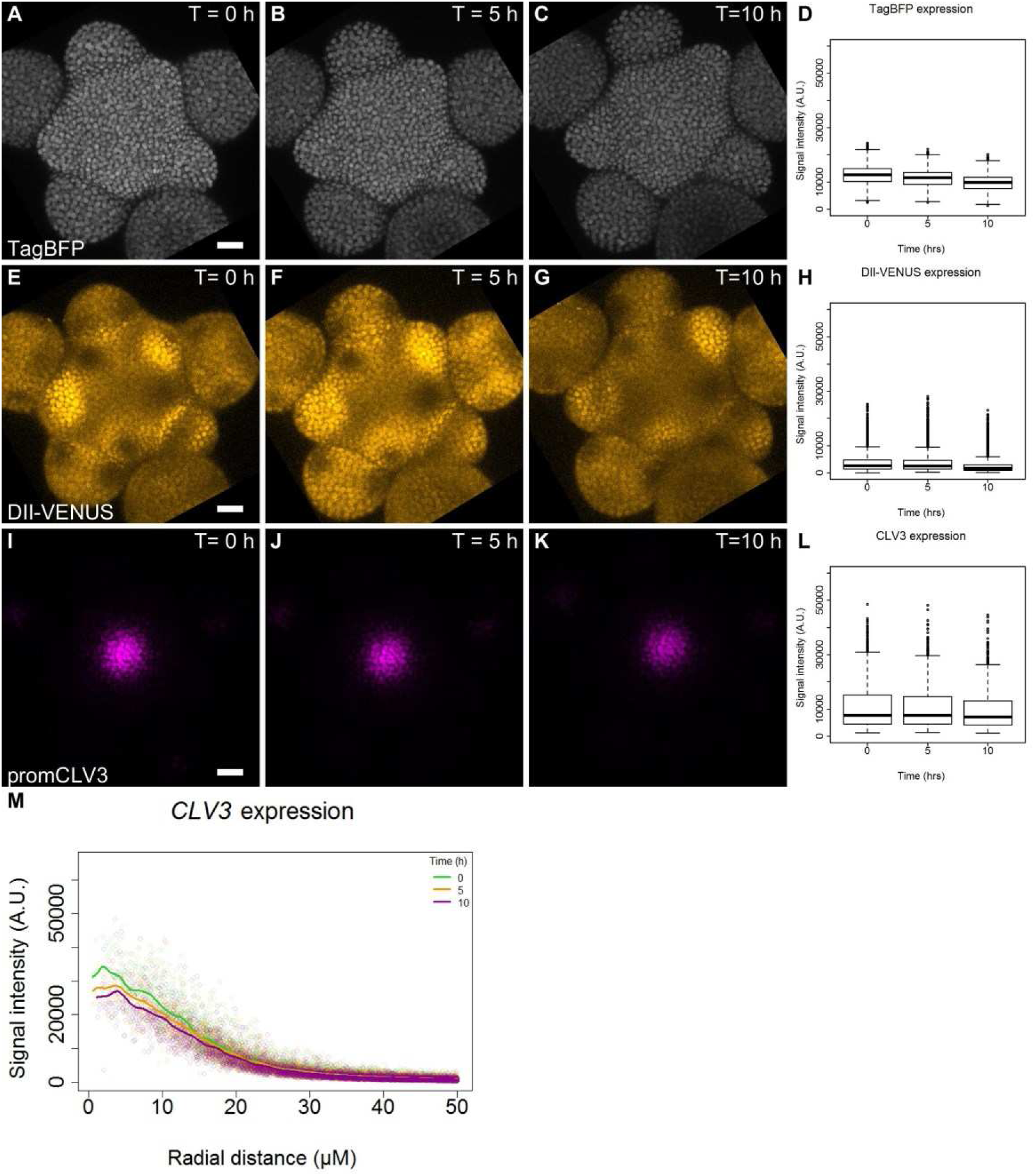
Expression pattern of qDII and CLV3. **A-C.** Representative images of RPS5a:qDII. TagBFP channel (grays). **D.** Quantification of TagBFP nuclei intensity at the L1 cell layer of the meristem (N=42991 nuclei). **E-G.** DII-VENUS expression pattern. **H.** Quantification of DII-VENUS-N7 nuclei intensity at the L1 cell layer. **I-K.** Expression pattern of pCLV3:mCherry. **I.** Quantification of CLV3 nuclei intensity at the L1 cell layer (N=6003 nuclei). Scale bars = 20 μM. **M.** Comparison of CLV3 expression as a function of radial distance from the center in time. Each point represents a nuclei and regression curves for each time point distribution. Boxplot centers show median.

**Figure S2.**
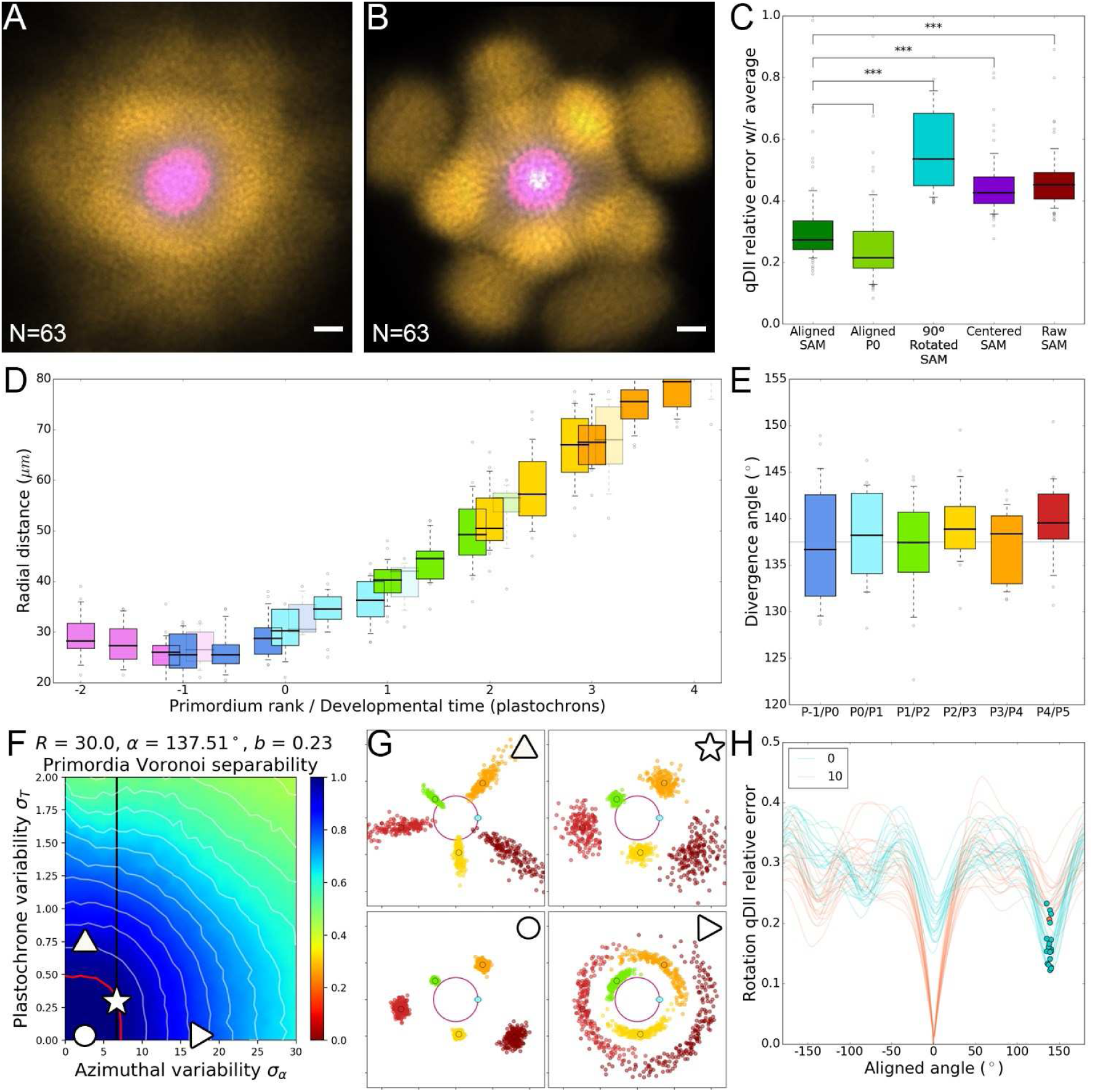
Precision of maxima positioning enables time extrapolation. **A.** Superposition of non-registered SAM images and registered by rotation. N=69 meristem images. **B.** Superposition of the same images in (A) after rigid registration. Scale bars = 20 μM. **C.** Distributions of relative map errors comparing an individual qDII map to the average of the rest of the population. From left to right, the boxplots represent the errors obtained using the aligned image positions as in **(B);** the aligned image positions but only in the Po domain; the aligned image positions compared with 90°-rotated individual maps; the raw image positions centered on the CLV3 domain; and the raw image positions as in (A). The alignment significantly reduces the error, not only in the P_0_ domain that is used for alignment but all over the SAM, making around 3 times less error that the worst possible alignment. **D.** Auxin maxima radial position as a function of developmental time, assuming a plastochrone time of 12h. Radial distance of an auxin maximum P_n_ at T= 0h lies between the previous maximum P_n-1_ at T=10h and T=14h. Boxplot centers show median. **E.** Relative divergence angle between 2 consecutive maxima. Boxplot centers show median. **F.** Theoretical study shows that variability vastly affects the separability of primordia clusters. In a phyllotactic model using the observed initial distance and speed (from **D**), divergence angle and angular variability (from **E**) parameter values the apparent separability of primordia across individuals can only be explained by very limited plastochrone variability (<0.3). Seamless superposition of individuals proves that the SAM achieves a very high rhythmic precision at population-scale. Red contour indicates 100% separability, white contours every lower 5*%.* Black line indicates experimental value of azimuthal variability. **G.** Primordia points obtained with a phyllotactic model for different values of angular and rhythmicity variability. Color marks the rank of the primordium. Symbols indicate the position in the parameter space (**F**) **H.** Computation of error between a map at T=10h and rotated maps (−180° to +180°) at T=10h (red), T=5h (green) or T= 0h (blue). Error values equal to 0 denote identical maps. The lowest error values with rotated maps (blue points) were globally found between the 10h map and the 0h map close to a 137° rotation, validating that the extrapolation heuristic of placing P_1_ at T=0h right after P_0_ at T=10h is indeed the most consistent way to extend a sequence in time

**Figure S3.**
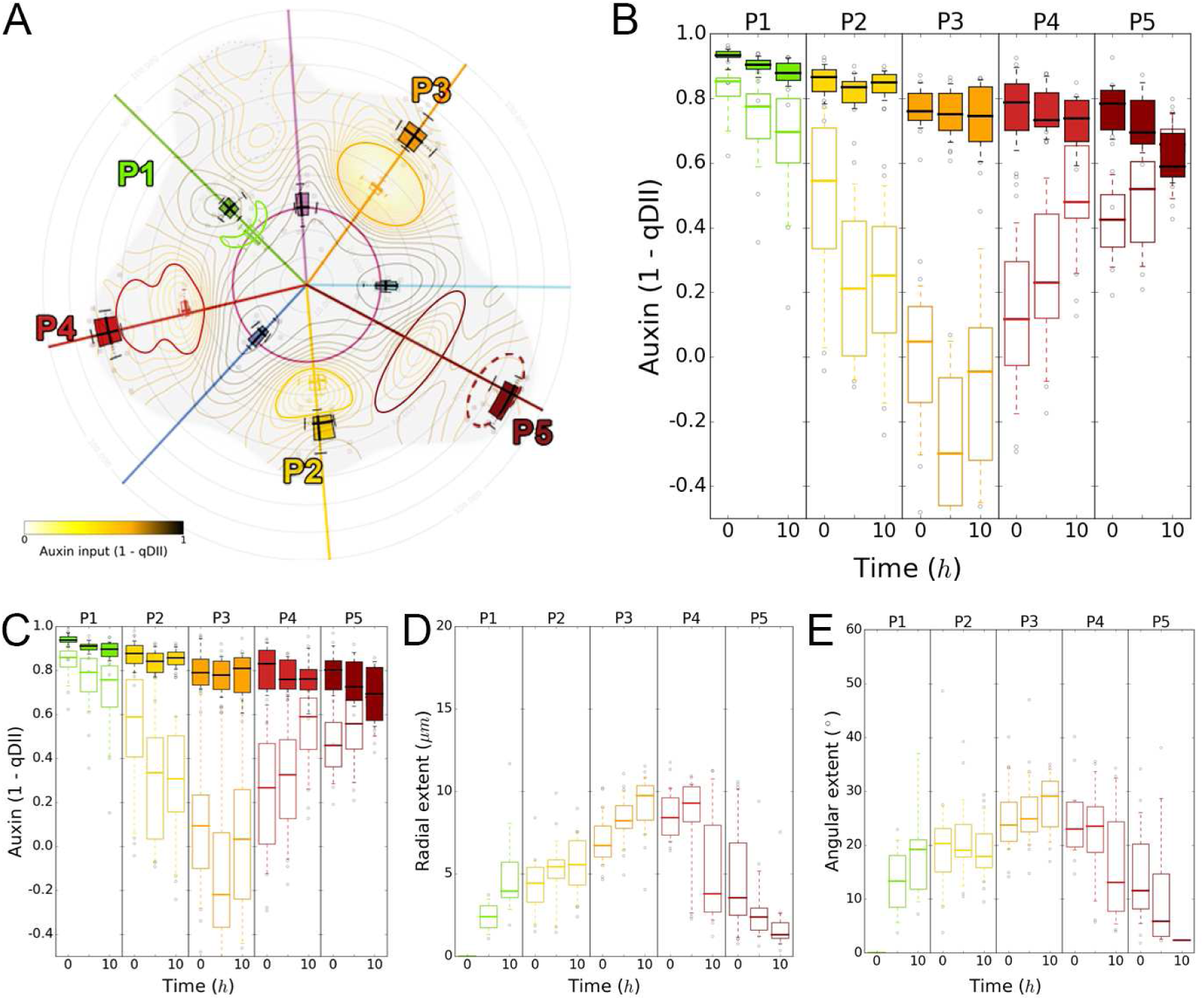
Auxin depletion areas are locally defined and transient in time. **A.** Auxin minima appear from stage P_1_ as a saddle point isolating radially the auxin maximum from the central zone, and progressively builds up until its reduction to a single cell file in the organ boundary at stage P_5_. **B.** Auxin values of maxima (filled boxes) and minima (outline boxes) in time for each primordia. **C.** Minima (outline boxes) closely follow their associated maximum (filled boxes) but stop at a fixed distance to the center after stage P_3_. **D.** Auxin depletion radial extend from the maxima to the center of the SAM. The depletion zone progressively widens in stages P1 to P4 before getting split in two at stage P_4_. **E.** Auxin depletion angular extent from the auxin maxima. Boxplot centers show median.

**Figure S4.**
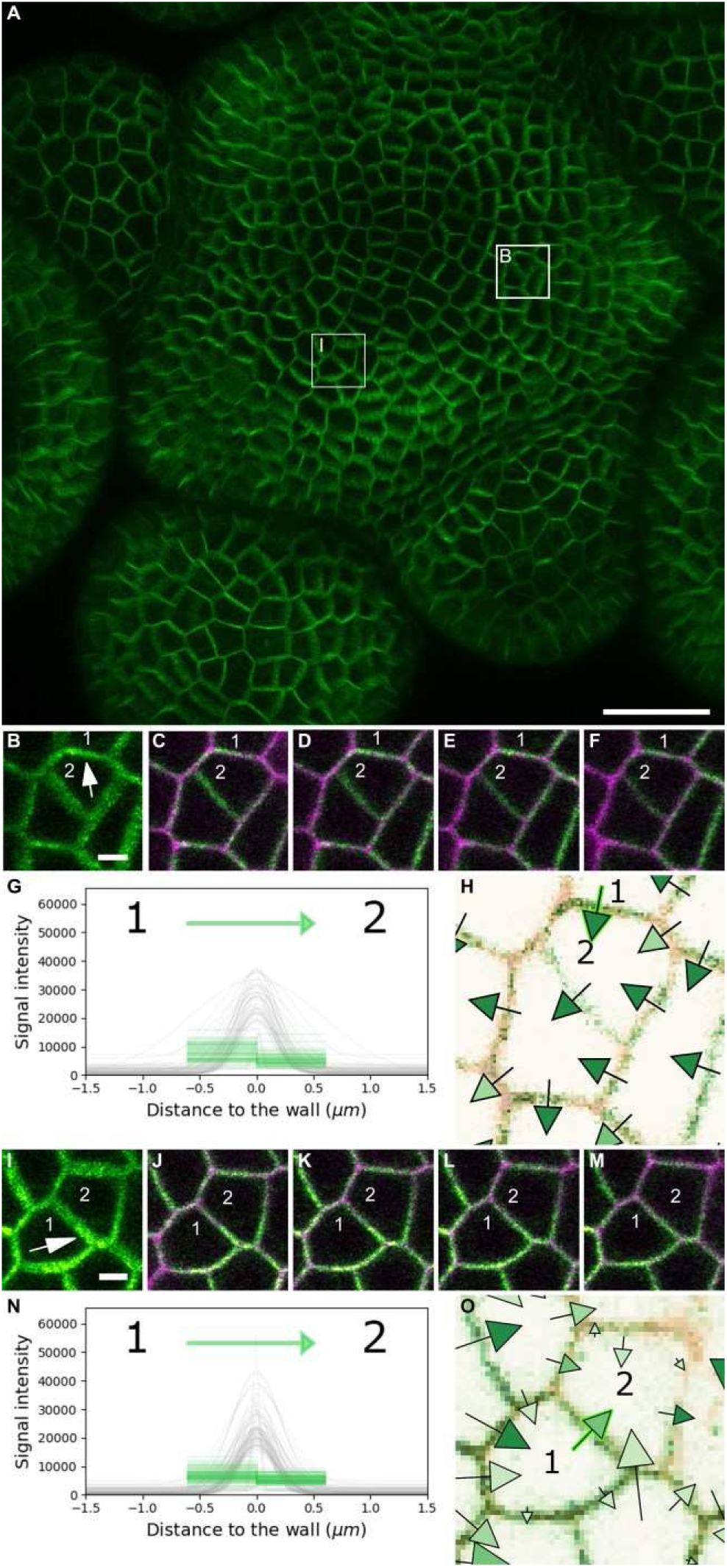
Croissant-shape PIN1 localization does not always reflect actual cell polarities. **A.** Confocal projection of PIN1-GFP. Magnified regions of the SAM in **(B)** and **(I)** are indicated by white squares. Scale bar = 20 μM. **B.** Confocal projection of PIN-signal. The arrow indicates the polarity for cell 2 assigned by visual impression based on intensity and croissant-shape of the signal. Scale bar = 2 μM. **C-F.** Z-slices of image in **(B)** with PIN 1 - GFP (green) and PI (magenta) signals. Notably the PIN 1 signal belongs to the membrane of cell 1 rather than cell 2 (y shift between signals). **G.** Average signal distribution for PIN 1 (green) and PI (grey) in the anticlinal wall between cell 1 and 2. Each line represents the average signal of a region of the anticlinal cell wall between cells. **H.** Polarity vectors detected by our method. Arrow with a green outline represents the polarity vector based on signal quantification in **(G). I.** Projection of PIN-signal. The arrow indicates cell 1 polarity assigned by visual impression as in **B.** Scale bars = 2 μM. **J-M.** Z-slices of image in (**I**) with PIN1-GFP (green) and PI (magenta) channels showing full overlap of both signals at the anticlinal cell wall between cell 1 and 2. **N**. Signal distribution for both signals in the wall between cell 1 and 2 as in **(G). O.** Polarity vectors detected. Arrow with a green outline represents the polarity vector based on signal quantification in (**I**).

**Figure S5.**
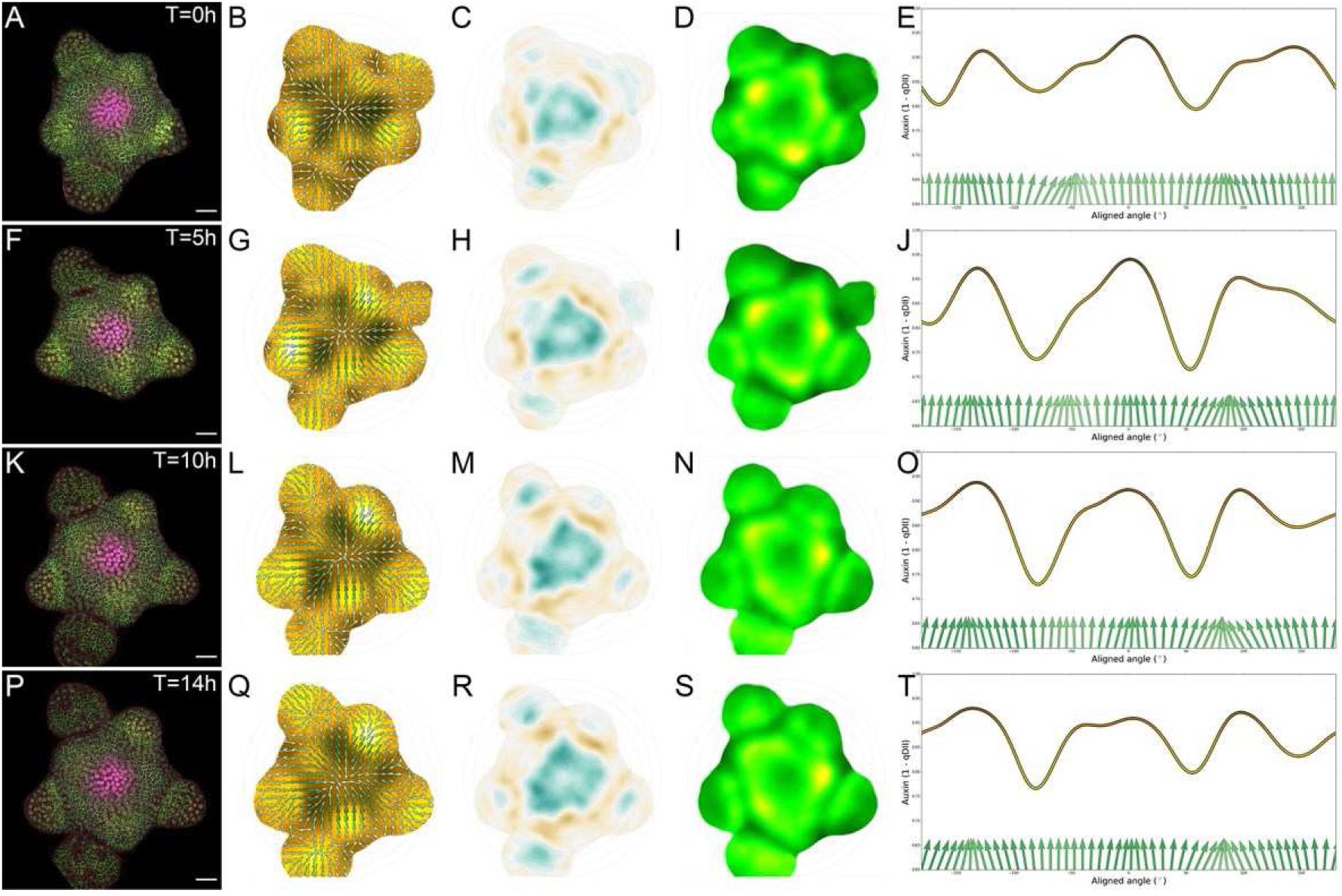
Vector fields and PIN global polarities Quasi-static PIN 1 network have a global convergence towards the center of the SAM. **A.** Representative confocal projection of the L1 cell layer of qDII (yellow), PIN1-GFP (green) and CLV3 (magenta). Average (N= 4 SAMS) auxin map and vector fields **(B)** showing convergence at the auxin maxima (black areas). **C.** local PIN1 convergence map. Blue represents convergence while brown represents divergent vectors. **D.** PIN 1 expression levels map. **E.** Circumferential auxin distribution at a 35 μ.M radial distance (line) with corresponding PIN1 vector fields (green arrows). **A-E.** T= 0 h. **F-J.** T= 5 h. **K-O.** T= 10 h. **P-T.** T= 14 h.

**Figure S6.**
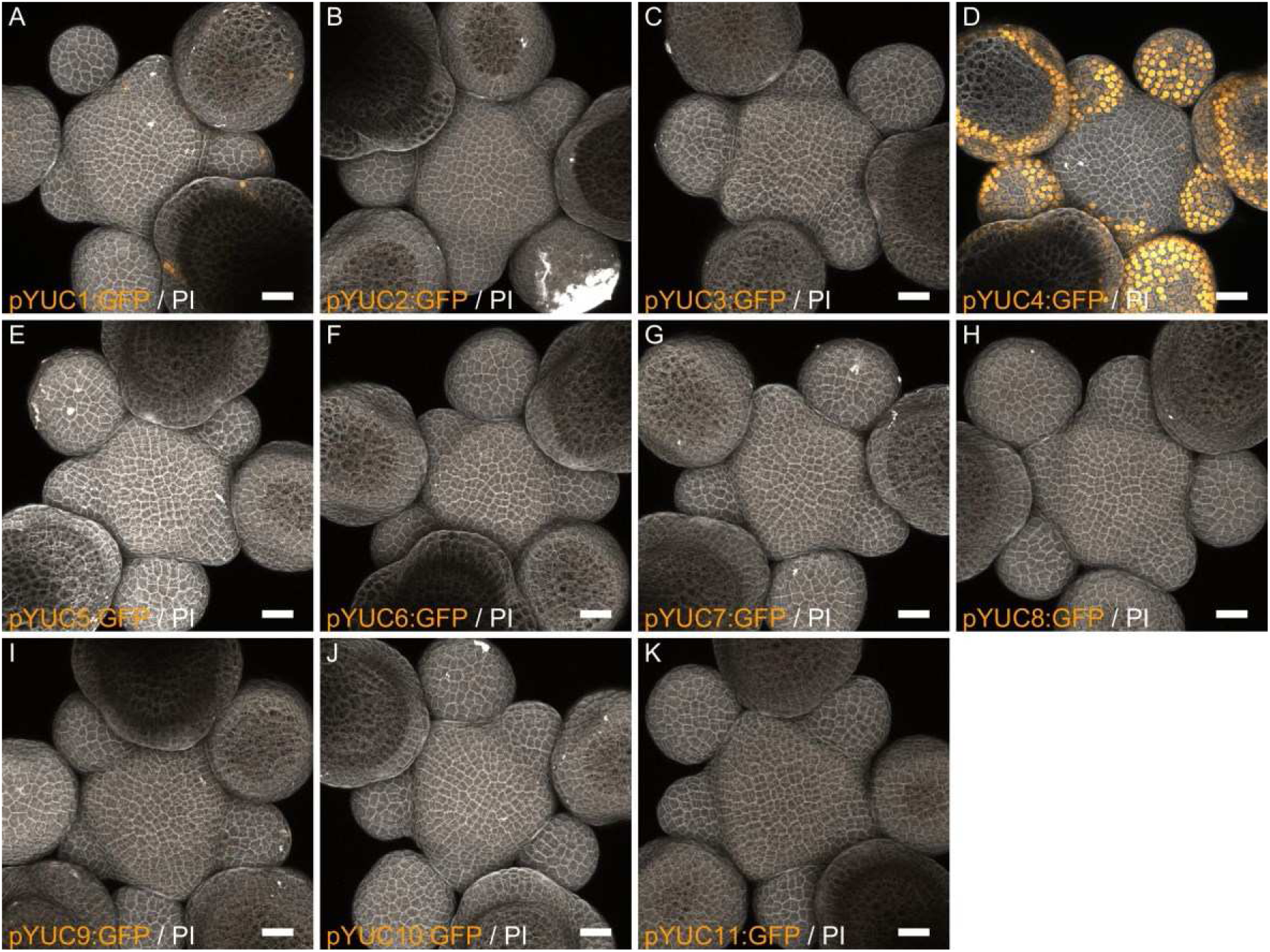
Expression patterns of YUCCA genes in the SAM A-K. Expression patterns of YUCCA genes in the SAM. Transcriptional reporter lines for the YUCCA auxin biosynthetic genes (orange) are indicated. Propidium iodide staining is shown in gray. Scale bars = 20 μM.

**Figure S7.**
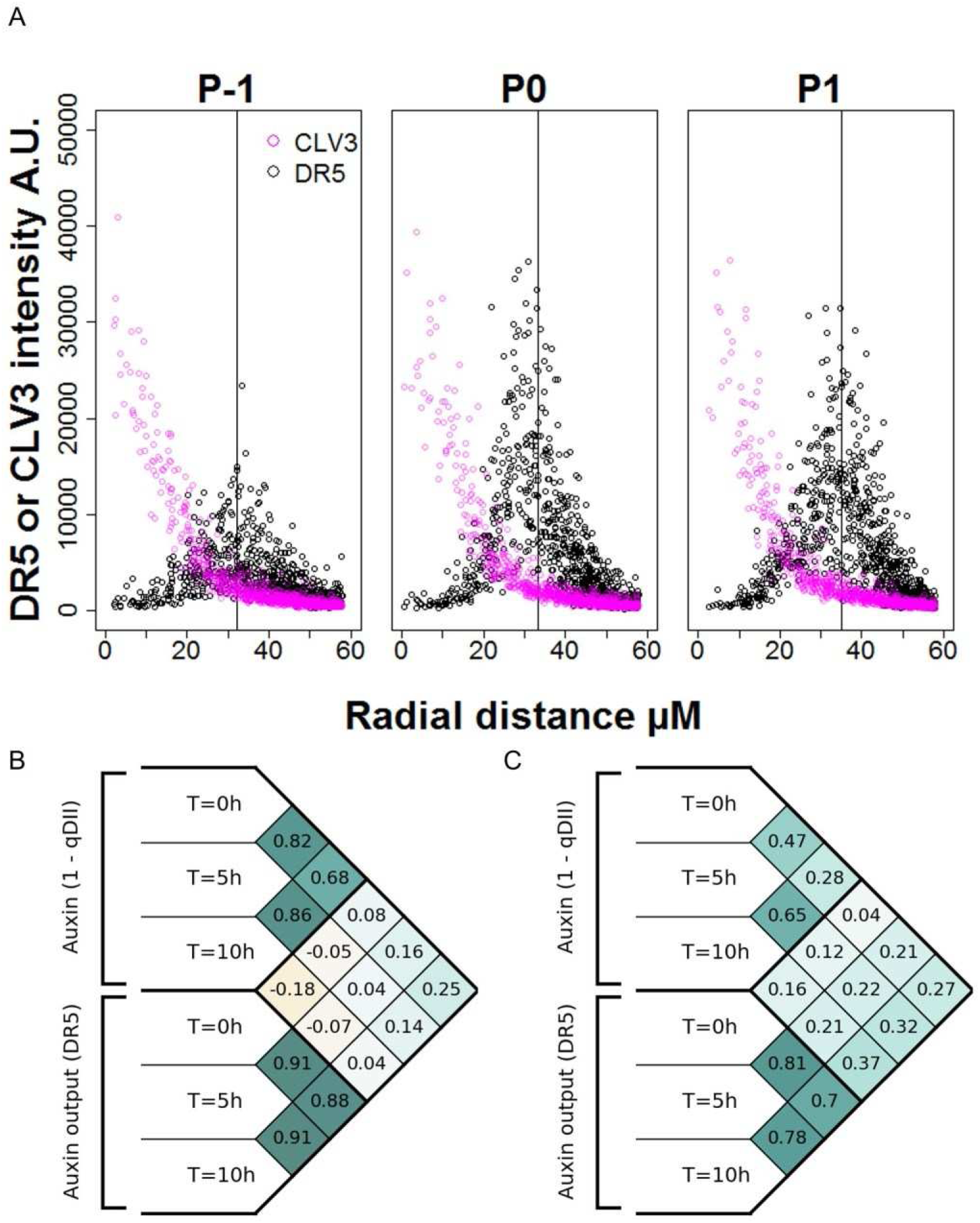
Precision of DR5 radial positioning and stage-dependent Pearson correlations. **A.** Radial distribution of CLV3 (magenta circles) and DR5 (black circles) at different locations of the SAM (P_-1_ to P_1_). N= 21 SAM. Each point represent a nuclei. Vertical line indicates the mean for DR5 radial distribution. **B-C.** Pearson correlation coefficients computed on tracked cell-level values of auxin input levels and output. The considered variables are the levels at time T=0h, T=5h and T=10h, the sum of auxin input from T=0h to T=5h and from T=0h to T=10h, and the variations of DR5 levels from T=0h to T=5h and from T=0h to T=10h. At a global level **(B),** no significant positive correlations are found between input and output variables, even when considering the time difference present in the data, arguing against a simple effect of delay, even between input levels and output variations, excluding a simple integrated response. Looking locally at nuclei in P_-1_ developmental stage makes stronger correlation values appear, and clear positive values (**C**). This overall suggests the existence of an exogenous stage-dependent variable controlling the relationship between auxin concentration and transcriptional response.

**Figure S8.**
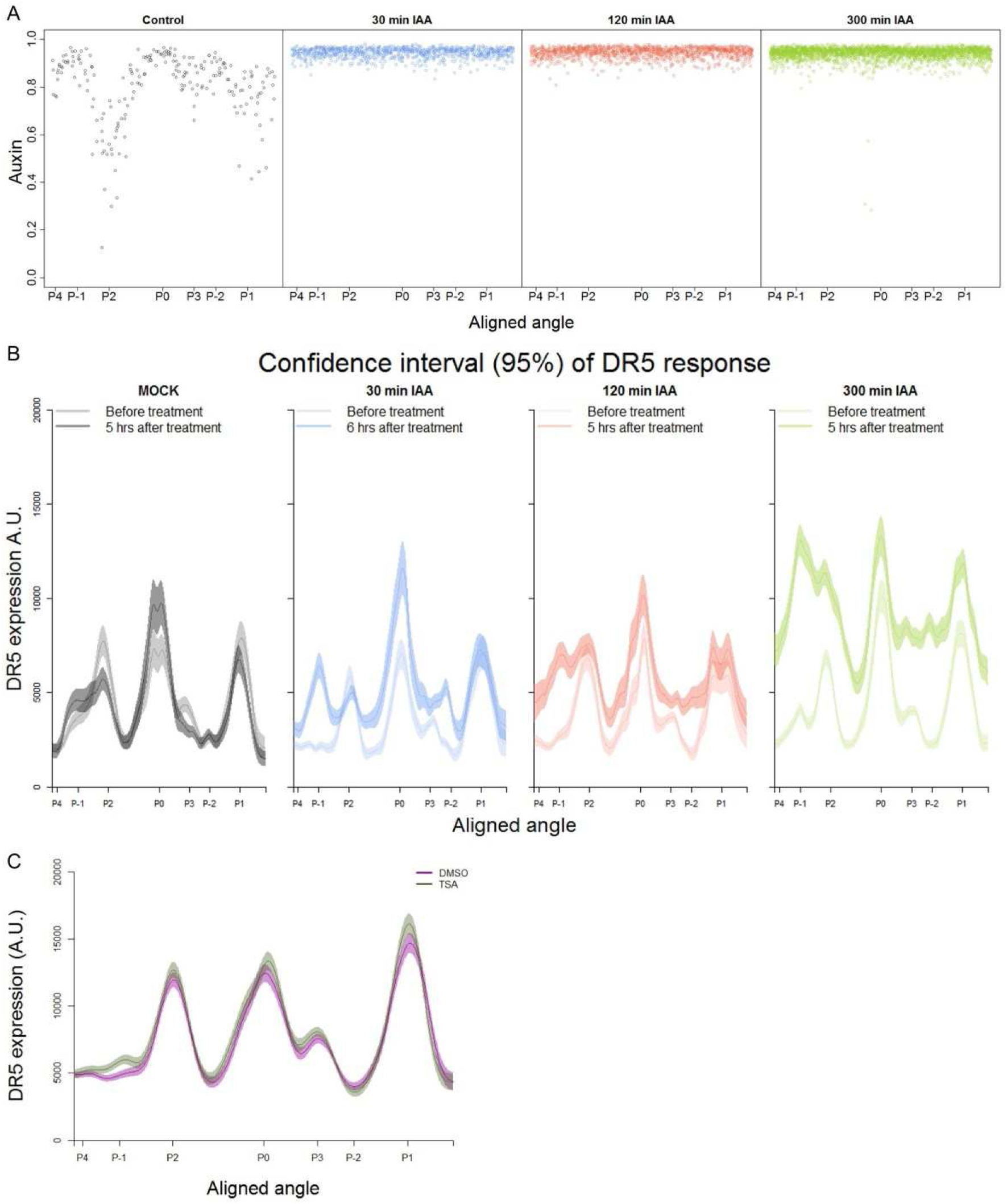
Time integrated auxin response is control by chromatin remodeling. **A.** Quantification of auxin (1-qDII) at the PZ after treatments. Each profile shows auxin levels after 1h for each treatment. Each dot represents a nuclei (Mock N=3045, 30 min N=1895, 120 min N=3378, 300 min N= 5081). The angular location of primordia are indicated on the abscissa. **B.** Confidence intervals (shade) and regression (line) for DR5 expression upon different auxin treatment lengths. Curves represent DR5 expression patterns at different regions of the PZ (ordinate) before and after treatment. Mock N= 1025 nuclei, 30 min N= 952 nuclei, 120 min N= 1344 nuclei, 300 min N= 2022 nuclei. **C.** Confidence Intervals and regression for DR5 expression of control (magenta) or TSA (green) treated meristems. DMSO N= 4184 nuclei, TSA N= 4141 nuclei.

**Table SI.**
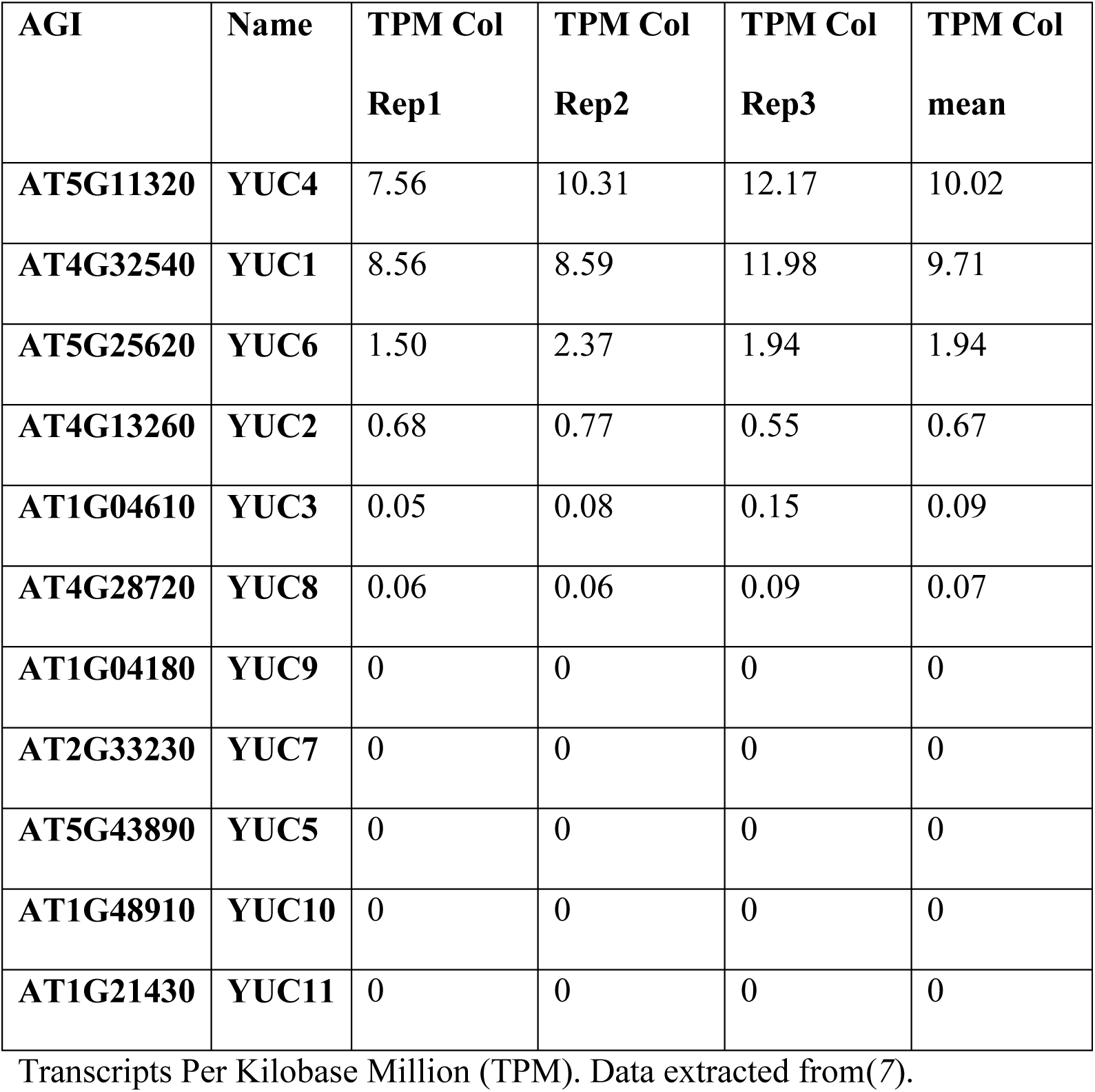
RNAseq expression of YUCCA genes at the SAM

**Table SII.**
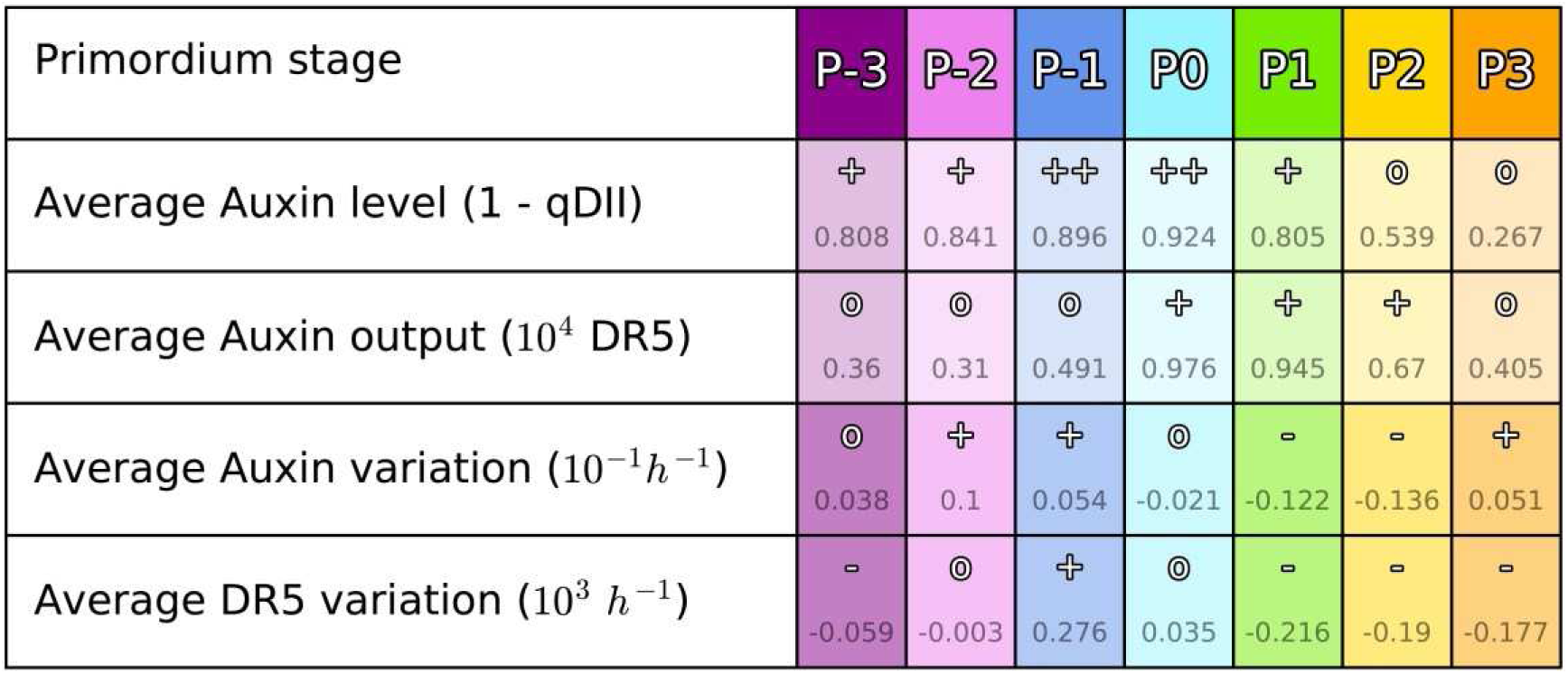
Auxin / DR5 variations in a primordium show a stage-characteristic behavior

Values of auxin and DR5 were quantified in tissue areas moving accordingly to cellular motion, across 10h of observation (rows 1 and 2) and their temporal derivatives estimated as the slope of a linear regression (rows 3 and 4). At the first order, each developmental stage from P_-3_ to P_5_ can be identified by a unique combination of the values of these four indicators. For instance a high stable value of both auxin and DR5 is characteristic of stage P_0_, whereas high increasing auxin with low increasing DR5 is characteristic of stage P_1_. “-”“0”,”+” and “++” correspond to arbitrary thresholds for the different variables: Auxin: “0” < 0.6 < “+” < 0.85 < DR5: “0”<5000<”+”; Auxin variation: ^M^-”<-0.05<”0”<0.05<”+”; DR5 variation: “-”<-100<”0”<100<”+”.

**Movie S1. Auxin developmental continuum over 9 plastochrons.**

Auxin (yellow to black) distribution dynamics in the SAM. P_0_ is located at the right.

**Supplementary Note 1. Definition of a conceptual frame for models and analysis**

The shoot apical meristem dynamically produces organ primordia, issuing from a central dome-shaped area, into a complex spatio-temporal pattern that is referred as phyllotaxis. In an abstract view of this structure, the meristem can be seen as dynamic collection of organ primordia characterized by their spatial trajectory relatively to the central zone (CZ) and by the evolution of their inner state. We propose a formal definition of such a system, which we name a “phyllotactic dynamical system”.

**Definition 1:** Let a **phyllotactic dynamical system *𝒮*** be a finite set of **primordia** considered on a time interval ***𝒯* =** [***t***_min_, ***t***_max_] ⊂ ℝ and such that:

- At every time *t* ∈ 𝒯, each primordium *p* ∈ *𝒮* is characterized by its current state {τ_p_ (*t*), *x*_*p*_ (*t*)-*y*_*p*_(*t*)} where:
  - τ*p* (*t*) G∈ [τ_min_, τ_max_] ⊂ ℝ is called the **developmental state** of the primordium.
  - *x*_*p*_ (*t*) = [*r*_*p*_(*t*), θ_*p*_(*t*), *z*_*p*_(*t*)] ∈ ℝ^3^ is the **spatial position** of the primordium in a cylindrical coordinate system, the origin of which is called the **center** of the system.
  - *y*_*p*_ (*t*) ∈ ℝ^*d*^ is a vector describing the **physiological state** of the primordium.
- The developmental state τ_*p*_ is a continuous strictly increasing function of time. Note that τ**_*p*_** is consequently a bijection between 𝒯 and τ_*P*_(𝒯).
- When it exists, the time 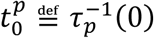 is called **initiation time** of the primordium *p.*
- The spatial position and the physiological state are conditioned by the developmental state of the primordium in such way that:

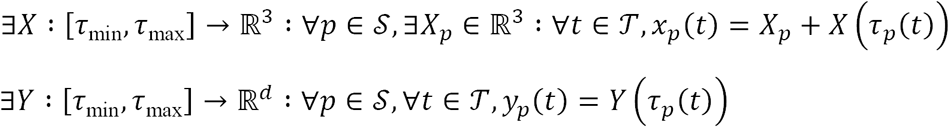
- *𝒮* is equipped with a strict total order denoted < that verifies:

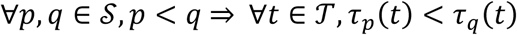

This definition reflects the idea that for any primordium, there exists an underlying physiological state, a hidden variable that determines all processes, both geometrical and physiological, that characterize primordium development. This state can be used to rank the different organs among them, and to run through the sequence of primordia in the order of their respective development. It is actually more common to refer to primordia by their integer rank in this developmental order:

**Property 1:** There exists a morphism between (*𝒮,* <) and (𝕫 <), and we can use it to denote the consecutiveness relationship in the strict total order of *𝒮* by:

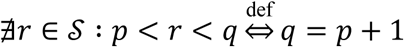

In a phyllotactic system, the notion of plastochrone refers to the time elapsed between two consecutive organ initiations. However it is common to speak about the plastochrone as a characteristic of the system when this duration does not vary over time:

**Definition 2:** We say that a system *𝒮* has a **plastochrone** *T* ∈ ℝ if two consecutive primordia in the strict total order of *𝒮* always have their initiation times separated by a time interval of length *T*:

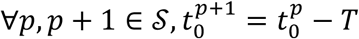

A stronger assumption that is generally made on a phyllotactic system is that it develops in a steady regime, meaning that it maintains a constant rate of development. This translates into linear functions for the developmental states of primordia with a common strictly positive slope. If we add the existence of a plastochron, then this slope is naturally equal to the inverse of the plastochron:

**Definition 3:** We say that a system *𝒮* with a plastochrone *T* has a **steady development** if all primordia in *𝒮* have their developmental states increasing at the same constant rate 1/*T*:

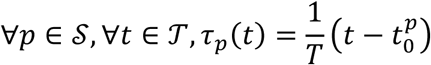

**Property 2:** In a system *𝒮* with a steady development of plastochrone *T*, all primordia increase their developmental state by 1 after a period *T:*

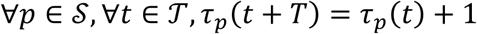

**Property 3:** In a system *𝒮* with a steady development of plastochrone *T,* two consecutive primordia in the strict total order of *𝒮* always have their developmental states separated by 1:

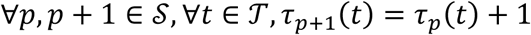

**Property 4:** In a system *𝒮* with a steady development of plastochrone *T,* at any time *t* ∈ 𝒯, the rounding function of the developmental state:

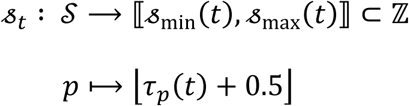

is an isomorphism. We call 𝓈_*t*_(*p*) the **developmental stage** of primordium *p* at time *t.* If 𝓈_*t*_(*p*) *= k* ∈ 𝕫 we say that the primordium *p* has the label ℙ_*k*_ at time *t.*

**Property 5:** In a system *𝒮* with a steady development of plastochrone *T,* two consecutive primordia in the strict total order of *𝒮* always have consecutive developmental stages:

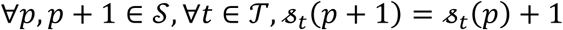

**Definition 6:** In a system *𝒮* with a steady development of plastochrone *T,* we can extend the notation of the consecutiveness relationship in the strict total order of *𝒮* using the isomorphism 𝓈_*t*_ to identify the primordia by their relative developmental stages write:

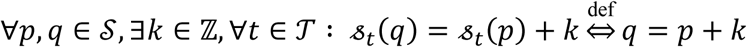

**Definition 4:** We say that a system *𝒮* with a steady development of plastochrone *T* has a **regular phyllotaxis** of divergence angle *α* if the constant parts of spatial positions of two consecutive primordia in the strict total order of *𝒮* only differ by a rotation of angle *α* around the vertical axis:

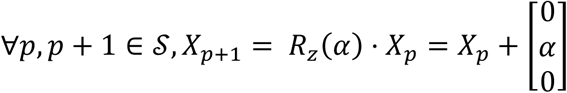

**Property 7: Spatio-temporal periodicity:** In a system *𝒮* with a steady development of plastochrone *T* and a regular phyllotaxis of divergence angle *α,* the system verifies the following spatio-temporal periodicity:

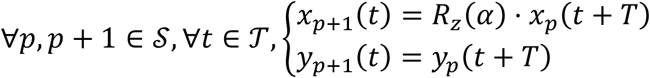

Phyllotaxis regularity offers a way to access the ranking of primordia simply by looking at their spatial positions. For instance, if the distance of a primordium to the center is increasing with time, then the order on distances reflects the order on primordia developmental states. Cues from distances and angular positions can typically be combined for an even more robust ranking of organ primordia.

**Property 8: Ordering on a regular phyllotaxis:** In a system *𝒮* with a steady development of plastochrone *T* and a regular phyllotaxis of divergence angle *α,* if *α* is not a simple fraction of 2π, i.e if:

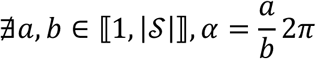

then the primordia angles θ_*p*_ are sufficient to know the strict total order on *𝒮*:

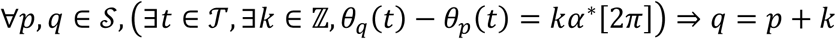

In this case, we say that *𝒮* has a clear regular phyllotaxis.

With this conceptual frame in mind, we can define thoroughly the problem addressed when labeling the primordia on a meristem observation, i.e. typically when one tries to estimate where is ℙ_1_ and where is ℙ_2_ on a microscopy image. In that problem, only a partial state is observed for each primordium, containing mostly its spatial position and possibly some quantified features. Using this information only, the goal is to stage the visible primordia, by affecting them a label that is as close as possible to their actual developmental state, in such way that if the method is used on different observations, the primordia assigned the same label really have close actual developmental states.

**Problem 1: Assignation of developmental stages:** Given a system ***𝒮,*** observed at ***n*** discrete time points **{t_1_,…,t_n_}** in which, for every ***p*** ∈ *𝒮* and for every ***t*_*i*_** ∈ **{*t*_1_, …, *t*_n_},** only a partially observed state 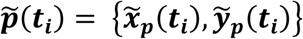 is available;

Find for every *t*_*t*_ ∈ {*t*_1_,…,*t*_n_} an estimated developmental stage function 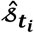 that verifies:

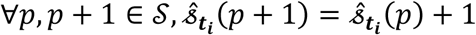

and that minimizes the average staging error of the primordia:

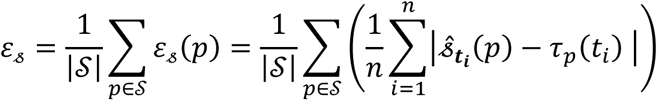

**Definition 5:** We say that a system *𝒮* is 𝕡 _*k*_ -maintaining (*k* ∈ 𝕫) if at all times, there is a primordium that has the label 𝕡_*k*_:

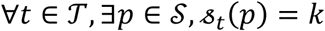

**Property 9: Reduced 𝕡_*k*_ assignation problem in a clear regular phyllotaxis:** Let *𝒮* be a 𝕡_*k*_ - mantaining system with a steady development of plastochrone *T* and a clear regular phyllotaxis of divergence angle *α*. The solution to the assignation of developmental stages problem can be reduced to:

Find for every *t*_*i*_ ∈ {*t*_1_,…, *t*_*n*_}, the primordium *p* ∈ *𝒮* such that 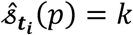.

**Definition 6:** Let *𝒮* be a system with a steady development of plastochrone *T.* We say that the physiological state function *Y* is 𝕡_*k*_ -characteristic (*k* ∈ *𝕫*) if there exists a value set *⌈*_*k*_ ⊂ #x211D;^*d*^ such that:

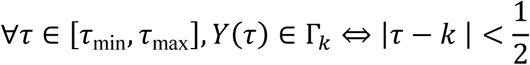

**Property 10: Developmental stage stability condition:** Let *𝒮* be a system with a steady development of plastochrone *T.* We say that a developmental stage assignation is stable if it is the same at all times of observation, i.e. if:

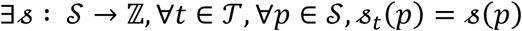

If *𝒮* is observed during a time interval smaller than its plastochron, then there exists a stable developmental stage assignation with a staging error bounded by 1/2 for every *p* ∈ *𝒮:*

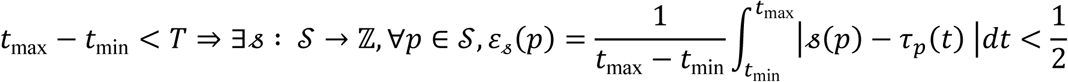

**Property 11: Stable 𝕡 _*k*_ -characterization solution:** Let *𝒮* be a 𝕡 _*k*_ -mantaining system with a steady development of plastochrone *T,* a clear regular phyllotaxis of divergence angle *α* and a 𝕡 _*k*_ -characteristic state function. If *𝒮* is observed during a time interval smaller than its plastochrone *T,* then the reduced stable solution to the assignation of developmental stages problem given by:

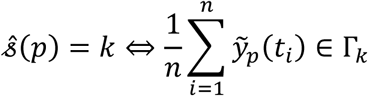

has a staging error bounded by 1 for every *p* ∈ *𝒮:*

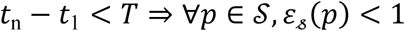

In a classical inhibitory field model of phyllotaxis, the developmental state τ_p_ = 0 of an organ primordium *p* corresponds to the time where an initiation is decided in the peripheral zone (PZ) and would therefore match a local spatio-temporal minimum of the global inhibition field. With the idea in mind that the depletion of auxin has very often been related to the concept of “inhibition” from those models of phyllotaxis, we consider that the instant where initiation happens corresponds to a local spatio-temporal maximum of auxin in the meristem. In other terms a characteristic of the 𝕡_0_ primordium should be that it has the maximal auxin level across the PZ.

Therefore if the local auxin maximality is the *j*^th^> component *Y*_*j*_ ∈{0,1} of the state function, then we consider that the function *Y* is 𝕡_0_-characteristic with Г_0_ = ℝ x … x]l/2,1] x … xℝ. We observed the meristems over a time interval of 10 hours, which is less that the estimated plastochrone in our experimental conditions. Therefore, by Property 11, we can define a stable assignation of developmental stages to the visible primordia of bounded error by assigning the label 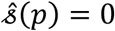 to the primordium that has the most often the maximal value of auxin across the PZ over the time of observation. If the meristems prove to be close enough to phyllotactic systems with a plastochrone and a regular phyllotaxis, then this first assignation will be enough to derive the developmental stages of all the other organ primordia based on their spatial positions. The method developed to perform this developmental stage assignation heuristic on experimental data is detailed in Supplementary Method 2. Evidence for the regularity of the observed phyllotactic systems is discussed in Supplementary Method 1.

### Supplementary Method 1. Effects of variability on a computational phyllotactic system

In this section we develop a formal study on regularity in a phyllotactic system. Notably, we wondered to which extent the apparent similarity of the observed SAMs could be informative on the level of precision in the process of organogenesis. To answer this, we simulated a sample of phyllotactic patterns assuming that i) they are all aligned with respect to the position of their 𝕡_0_ ii) their plastochrones and divergence angles are drawn from random distributions centered on a common average value. By varying the levels of noise on both angular positions and plastochrones, we assess how variability impacts the overlapping of phyllotactic patterns at the scale of a population.

Let us consider a 2D phyllotactic dynamical system *𝒮* (see Supplementary Note 1) formed by *N* organ primordia observed on a temporal interval *𝒯.* At every time *t ∈ 𝒯,* each primordium labeled *p* is represented by its developmental age τ_*p*_ and by its coordinates in a 2D cylindrical reference frame:

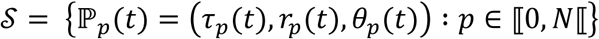

If we assume that the system has a plastochrone *T* and has a steady development, all primordia develop at the same constant rate 1/*T.* In that case, we can derive:

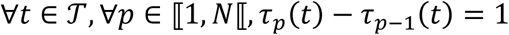

In addition, if we consider that the system has a regular phyllotaxis with a divergence angle α, a central zone radius *R,* and an exponential radial motion law of coefficient *β* we can write the state of the system as follows:

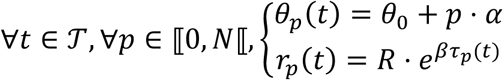

This can be translated into incremental equations to obtain a recursive definition of the state of the system, knowing 𝕡_*0*_(*t*):

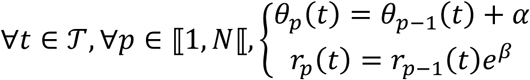

We set ourselves in a context where all the considered phyllotactic systems have previously been aligned on 𝕡_0_, i.e. where ∀*t* ∈ 𝒯,*θ*_0_(*t*) = 0 and τ_p_(*t*) ∈ [–0.5,0.5].

We study what happens if we introduce variability into this system, by adding noise on two of the key variables of the system:

- A Gaussian noise of standard deviation *σ*_*α*_ on the divergence angle *α*
- A Gaussian noise of normalized standard deviation *α*_T_ on the plastochrone time *T*

To be more precise, we consider that the system still has a plastochrone and a constant development rate 1 /*T,* but that the gap between the initiation times 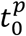 and 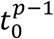 of two consecutive primordia that should always be equal to *T* is actually a random variable:

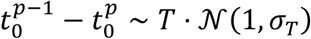

which translates into:

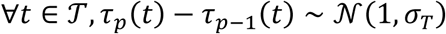

Consequently, we can formulate the recursive definition of the system as the drawing of *2*(*N —* 1) random variables:

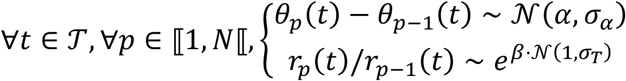

We simulate a population of such systems by generating *K* single-time instances that are all aligned on 𝕡_0_. To do so, we draw for each instance a value for τ_0_ from a uniform distribution in [—0.5,0.5], then use the initial values 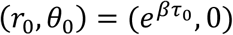 and construct the system recursively by drawing the corresponding random variables. This way, we obtain a population of organ primordia positions identified by their rank *p* (Fig. S9A).

In this random population, we are interested in which extent the generated phyllotactic patterns overlap. To do so, we estimate whether the points corresponding to primordia of the same rank can be grouped into separable clusters. Therefore, we consider the obtained primordia as a point cloud labeled by primordium rank:

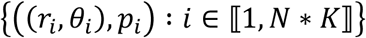

To answer the separability question, we measure to which extent the identically labeled points {(*r*_*i*_,θ_*i*_): P_*i*_ = p}p∈⟦0,*N*⟦ are separable by applying an unsupervised clustering algorithm. We chose to use the k-means method because it outputs a Voronoi diagram of the obtained centroid points, which places linear boundaries between clusters, and therefore measures linear separability of the point cloud, in an unsupervised manner. Prior knowledge was fed to the algorithm by setting the number of components to *N* and by initializing the centroids on the positions of the primordia in a model without any noise:

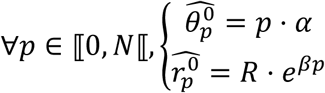

After convergence, the algorithm returns *N* centroid points 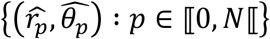 that we use to construct the Voronoi diagram. This actually defines a predictor for primordium rank by looking inside which cell of the diagram lies a given point (Fig. S9B):

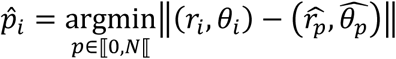

Finally, we estimate the Voronoi separability *v* of our cloud of primordia points by computing the accuracy of the primordium rank prediction (noting δ_*i j*_ the Kronecker delta on integers):

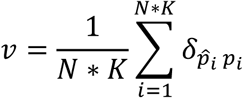

If *v* equals to 1, it means that the primordium points group into perfectly identifiable clusters. This can be interpreted as the fact that *K* randomly sampled individuals can be superimposed perfectly, once they have been centered and aligned on their primordium which is the closest to the 𝕡_0_ stage.

This will obviously be the case (as long as the motion coefficient β remains realistic, typically <1) if no noise is introduced into the system. If *σ* _*α*_ = *σ*_*T*_ = 0, then all individuals proceed from the same regular exact pattern, and the only variability will be the one of the instant of sampling τ_0_. We wondered up to which level of noise this separability property could be maintained, in order to understand what a high observed separability could tell us on the intrinsic regularity of a phyllotactic system.

To do so, we scanned the parameter space by varying *σ* _*α*_ between 0° and 20° and *σ*_*T*_ between 0 and 2 plastochrones, in a first time with the values *R* = 30*μm, α=* 137.51°, *β*= 0.23 corresponding to actual measured data (Fig. S2D-E). As expected, increasing the angular variability creates more elongated clusters (Fig. S9C) that still appear separated. Yet, the Voronoi separation introduces confusion between neighboring primordia, more specifically making 𝕡_*p*_ and 𝕡_*p*+3_ overlap (Fig. S9F). Interestingly, when we increase the plastochrone variability (Fig. S9D), the confusion concerns rather 𝕡_*p*_ and 𝕡_*p + 5*_ (Fig. S9G). In both illustrated cases, the separability score drops markedly below 95%, while the separability of the actual observed data has been evaluated in the same way at 100% (Fig. 9E).

The landscape of separability in the *σ* _*α*_ x *σ*_*T*_ parameter space gives an insight on the effects of variability on a population of individuals (Fig. S2F-G). With no surprise, primordia points appear to be less and less identifiable as azimuthal or plastochrone variability increase, and even worse when both do. But it shows that there exists a maximal level in variability up to which the clusters are still perfectly separable (Fig. S2F, red contour). Interestingly, the azimuthal variability we measure on our observed SAMs is close to the upper bound of *σ*_*α*_ on this optimal subspace.

If we fix the value of *σ*_*α*_ to 6.7° as measured in our data (Fig. S2F, black vertical line), then the maximal plastochrone variability that is allowed for the separability to remain at 100% is close to 0.3. This means that it would be impossible to see the near-perfect superposition observed in our data if the phyllotactic system that produced it had a plastochrone variability greater than 0.3, which would translate into an uncertainty on organ initiation times of nearly 3 hours. We have therefore demonstrated that the SAM achieves, at least, this level of rhythmic precision (though it might even be more precise) at the scale of a population (*N=*21). This consequently validates our first order assumption that all considered SAMs are in a steady regime of development with the same plastochrone duration, and that the whole set of individual primordia of a given rank forms a homogeneous population in terms of developmental age.

Another interesting feature evidenced by this analysis is the influence of the motion speed of primordia separability. We explored the *β* x *σ*_*T*_ parameter space by fixing the value of *σ* _*β*_ to the observed one, and varying the motion coefficient *β* between 0 and 0.4 (Fig. S9H). It appears that lowering the speed reduces the maximal possible value of *σ*_*α*_ to achieve 100% separability, as the points tend to overlap more in the radial dimension, leading to a decreasing separability at fixed angular variability. On the other hand, increasing the speed seems to greatly affect the tolerance to plastochrone variability. For instance a value of *σ*_*T*_ = 1 that translates into a separability of 95% when *β =* 0.1 suddenly drops to a separability of only 50% if *β* is increased up to 0.4. However increasing motion speed does not affect this much the maximal value of *σ*_*T*_ required to achieve 100% separability, which always remains close to 0.3. This consolidates our previous conclusions, even in the case of an underestimation of the radial speed.

**Figure S9.**
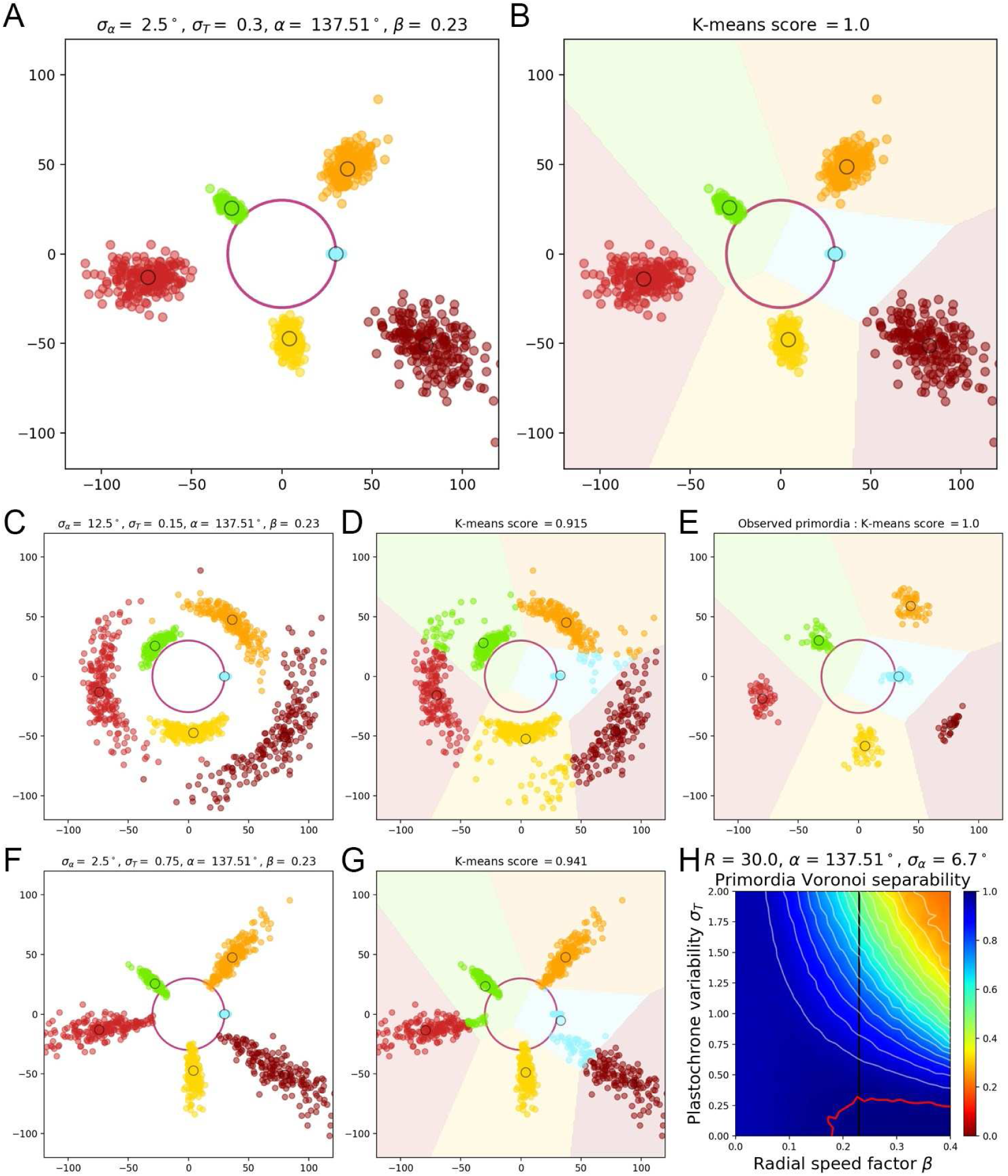
Primordium separability requires limited angular variability and highly precise rhythmicity. **A.** Primordium points are generated from a computational phyllotactic model with a control over the variability in angular positioning and plastochrone duration. **B.** Linearly separable clusters in the resulting point cloud are identified using an unsupervised algorithm with prior information. The obtained labeling in primordia ranks is compared with the theoretical one to compute a separability measure. **C-D.** Increasing the azimuthal variability creates clusters that are more difficult to separate and generates confusion between P_n_ and P_n+3_. E. The primordia point from the observed experimental data form perfectly separable clusters F-G. Increasing the plastochrone variability creates clusters that are more difficult to separate and generates confusion between P_n_ and P_n+5_. **H.** Separability evaluated by varying speed coefficient and plastochrone variability. Modifying the radial speed of primordia changes the tolerance of the system to azimuthal and plastochrone variability. High rhythmic precision is always required to achieve seamless superposition. Red contour indicates 100% separability, white contours every lower 5%. Black line indicates experimental value of speed coefficient.

**Figure S10.**
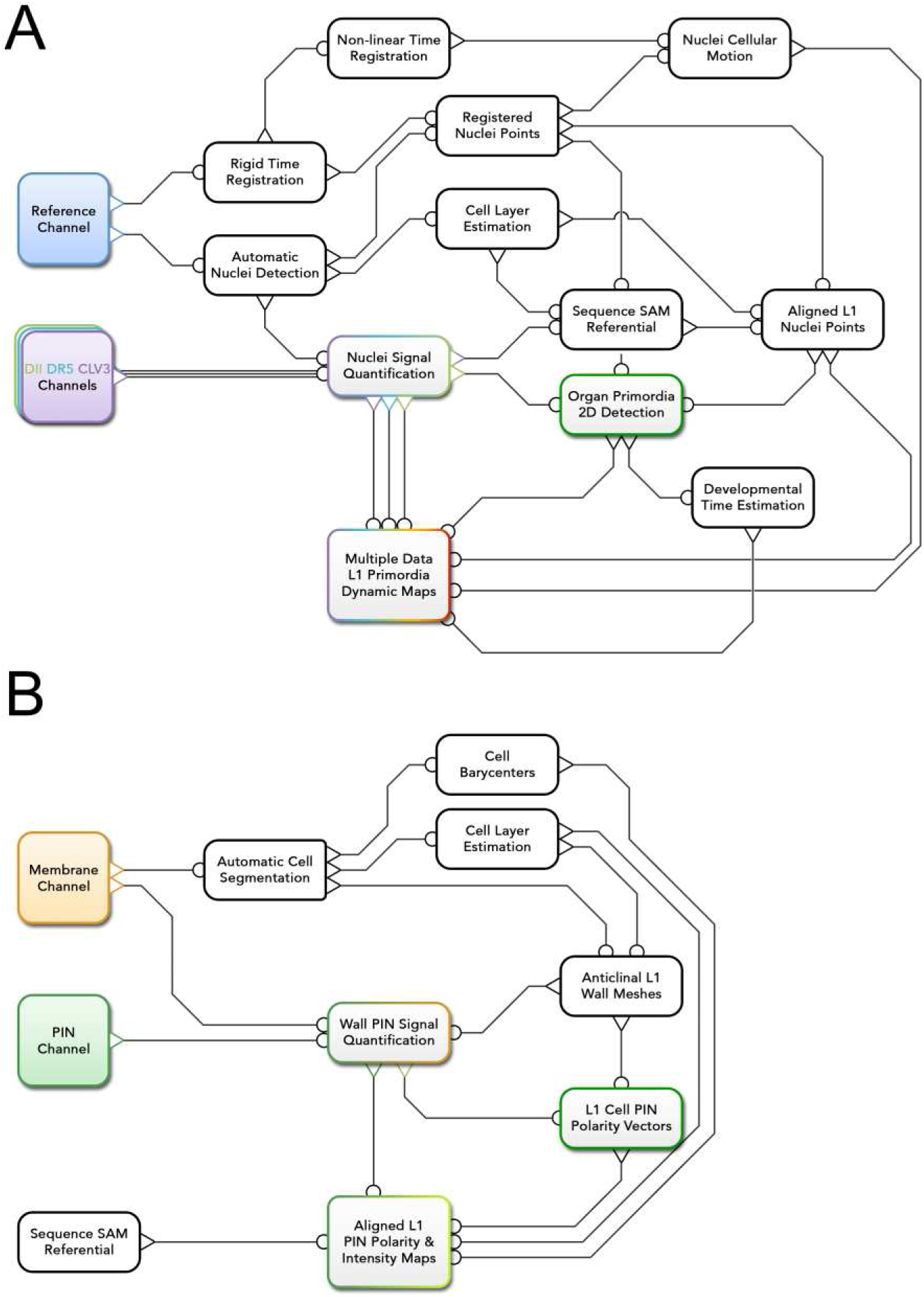
Automatic quantification pipelines of time-lapse images. **A.** To obtain quantitative data from the images produced under the microscope, various sequential processing steps need to be performed, from the extraction of the relevant objects (nuclei positions with their different channel intensity values) to the geometrical characterization and the spatio-temporal registration of the tissues, to finally get a complete, aligned and consistent dataset gathering all the imaged meristems. B. The quantitative estimation of PIN polarity relies on the analysis of cell wall and membrane-marker images that need to be processed differently. An alternative automatic pipeline performs the necessary steps, from the segmentation of the cells and the extraction of L1 anticlinal walls to the quantification of signal distribution at each cell wall and the reconstruction of the polarized cell network.

### Supplementary Method 2. Computational pipelines for image and data analysis

### 1. Nuclei marker image quantification and aligned L1 signal maps

Confocal microscopy images saved as CZI files are processed using a complex computational pipeline that involves several image analysis, computational geometry and data manipulation steps, performed sequentially as depicted in Fig. S10A.

#### Image reading

CZI files produced through the ZEN software(*8*) of the LSM-710 microscope are opened using a Python script(*9*) and spL1t into independent channels that are saved separately as INR image files(*10*). This operation preserves all the information contained in the raw image format. In the specific case of acquisitions for which both *pPINl.PINl-GFP* and *pRPS5a:DII-VENUS* are imaged, the close emission wavelengths causes the PIN1 -stained cell membranes to appear in the DII nuclei images. In that only case, the PIN1 signal intensity is subtracted from the DII image channel before saving the file.

#### ROI cropping

We use the polygonal selection tool of the ImageJ software to manually define a region of interest in all the slices of every image, and doing so, digitally dissect the outermost organs to get meristem images with at most 6 visible organs. This binary mask is then applied on all the channels, and masked channels are saved in separate INR files.

#### Automatic nuclei detection

For each image, we perform an automatic detection of nuclei points based on the masked *pRPS5a:TagBFP* channel only. It consists of a 3D image stack 𝒥 _Tag_ defined on a regularly spaced voxel grid {𝒥 _Tag_(*x, y, z*): (*x,y,z*) ∈ *I*} that has a potentially different voxel size *v* on each dimension so that:

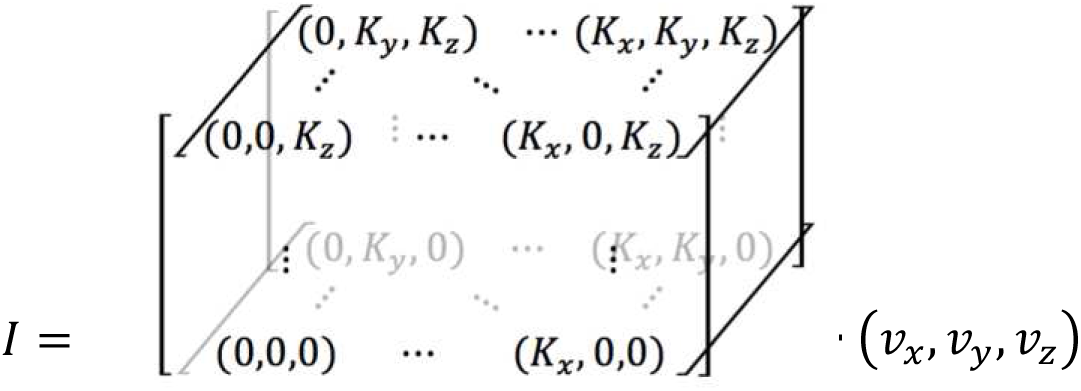

The image is converted into a 4D Gaussian scale space(*11*) by filtering it successively with *K*_*σ*_ 3D isotropic Gaussian filters 𝒢 of increasing radii, varying geometrically between *σ*_*min*_ and *σ*_*max*_:

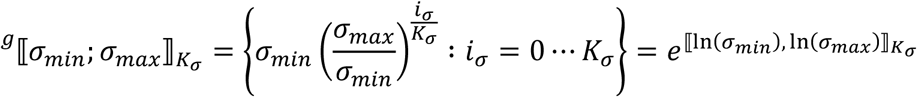

Nuclei are detected as local 4D maxima in this scale space representation noted 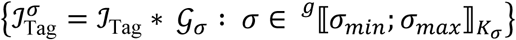, with the L1mit that only the maxima where the response 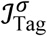 is higher than a threshold *𝒥*_*min*_ are retained. A 4D point (*x,y,z, σ*) is then considered a local maximum if and only if:

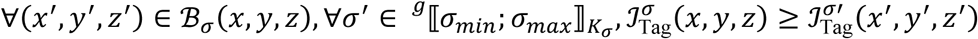

where ℬ_*σ*_ (*x, y,* z)is the discrete ball of radius *σ* in the image voxel grid:

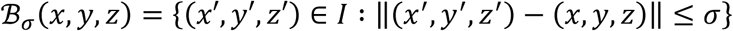

The detection results in a 3D point cloud 𝒩 where each detected nuclei *n* ∈ 𝒩 is represented by a single position *P*(*n*) = (*x,y,z*) in physical coordinates. This detection method has been evaluated on a set of 4 manually expertized SAM images acquired at different voxel sizes with a 16 bit encoding. The parameter testing led to the determination of the optimal values *K* _*σ*_ = 3, *σ* _*min*_ = 0.8*μm, σ* _*max*_ *=* 1.4*μ m, 𝒥*_*min*_ = 3000 corresponding to an evaluated performance of 95.6% recall and 98.5% precision.

#### Nuclei signal quantification

Every detected nucleus is assigned one value of signal intensity per image channel. This value is obtained by computing a weighted average of the masked channel intensity *𝒥*_*S*_ around the position of the nucleus. The signal images showing some local subcellular noise, the raw voxel value might not be fully representative of the whole nucleus. We chose to use a distance-based Gaussian weight, of constant radius *σ*_*S*_ *= 2*μ*m* for all channels, to account for as much as possible of the signal information inside the nuclei, for which the typical measured diameter is ∼ 5 μ*m.* Practically, we first filter the signal image by a Gaussian kernel of radius *σ*_*S*_ and read the values at the voxel positions of all detected nuclei:

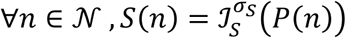

#### Cell layer estimation

For the purpose of the analysis, we want to discriminate between the first layer of cells (L1) and the rest of the tissue. We use an automatic method to do so, which in our case cannot rely on adjacency to the background as one would do on a segmented membrane-marker image. Instead we will use the distance of the nuclei to the estimated surface of the tissue. This surface is computed as a 3D triangular mesh based on the *pRPSa:TagBFP* channel. The image is filtered by a large Gaussian kernel to diffuse nuclei intensity between cells, and is thresholded to obtain a binary region, which is meshed by applying a Marching Cubes algorithm(*12*) on a resampled version of the image. This mesh undergoes a phase of triangle decimation(*13*) and isotropic remesbing(*14*) to obtain a surface composed of roughly 50000 regular faces. After normal estimation, only the largest connected component of triangles facing towards the upper side of the meristem is kept and used as the estimated meristem surface. For each vertex *v* of this surface mesh ℳ, we look for the closest detected nucleus by computing the distance to all of the points {*P*(*n*): *n* ∈ 𝒩}. The nuclei that we label as “L1” are the set of nuclei points that are closer than any other to at least one vertex of the surface:

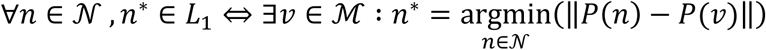

#### Rigid time registration

The previous steps were performed individually on each frame of the time-lapse acquisitions. To study consistently the dynamics at the scale of the sequence of 𝒯 images, we need to place the quantitative nuclei information in the same spatial reference frame. To do this, we estimate 3D rigid transformations between consecutive time frames of the sequence. This estimation is performed using a block matching algorithm(*15*) applied once again on the *pRPS5a:TagBFP* channel of the consecutive images. This produces 𝒯 –1 isometry matrices in homogeneous coordinates 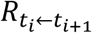 that can be inverted and/or multiplied to transform any frame of the sequence into the spatial reference frame of any other.

We use these matrices to transform the images into the reference frame of the first frame of the sequence (*t*_0_ = 0*h*). The transformed images are saved as separate INR files.

#### Registered nuclei points

We also apply the obtained transforms to the positions of the nuclei detected at time *t*_*i*_ to obtain their registered positions *P*^0^ in the reference frame of the first frame of the sequence:

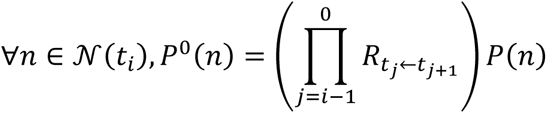

#### Non-L1near time registration

In a second time, we want to estimate the local deformation of the tissue, notably to get a quantitative measure of individual cell motion and to approximate local cellular growth. We compute this new transformation as a dense vector field that maps two consecutive rigidly registered *pRPS5a:TagBFP* images, using again the block matching framework. This approach has proven to be efficient on plant tissues in the case of L1mited deformations(*16, 17*) which is definitely the case for the meristematic tissue we are considering with a 4*h* to 5*h* interval between frames.

We do not use directly the resulting registered images but rather the vector field estimated in one of the two possible ways 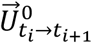 (respectively 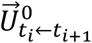) that stores at each voxel position a 3D vector measuring the local total deformation of the tissue to go from the current frame to the next one (respectively from the next frame to the current one). These two vector fields are saved as three-dimensional INR files.

#### Nuclei cellular motion

We estimate cellular motion between two consecutive sequence frames taken at times *t*_*t*_ and *t*_*i+*1_ in the forward direction using the first layer of registered nuclei points 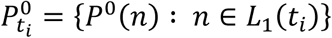 and the transformation 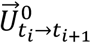 that maps the current frame into the next one by a vector field. Each nucleus can then be assigned a local motion vector 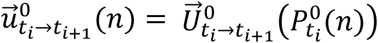

We deduce speed vectors measuring the local forward speed of cellular motion by dividing the motion vector by the time interval:

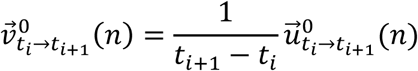

Conversely, similar cellular vectors are computed in the backward direction using the 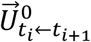 vector fields.

#### 2D maps of epidermal signal

We use a generic tool to take a signal defined on a discrete set of points and make it continuous in space. It basically computes a weighted average of signal values at every possible location in space. The weighting function we use is a parametric sigmoid density function η_*R,k*_ of the distance *r* to a point, which relies on two parameters, an extent parameter *R* and a sharpness *k* so that η_*R,k*_(0) ∼ 1, *η*_*R,k*_(*R*) = ½ and 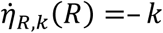 (Fig S11A) and takes the form:

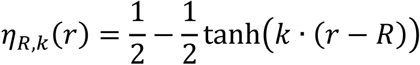

The continuous signal is then defined at every point *p* of space based on a point cloud {*P*(*n*):*n* ∈ 𝒩} and the associated signal values {𝒮 (*n*): *n* ∈ 𝒩} as:

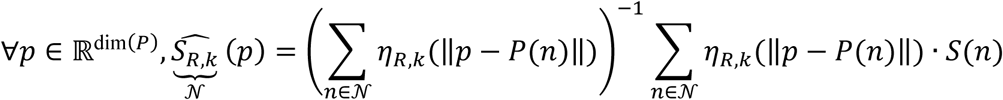

Note that, for any point *p* where ∑_*n∈N*_η_*R,K*_(‖*p–P(n)*‖) = 0, the signal map is not defined. To make the map outlines closer to their actual support, we will even consider that 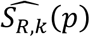 is defined only if ∑_*n∈N*_η_*R,K*_(‖*p–P(n)*‖) = ½, This constraint is equivalent to consider the implicit surface (or curve) obtained with the point cloud as generator and *η*_*R,k*_ as potential as the boundary of the definition domain of 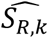.

In the rest of the analysis, we will mostly compute 2D maps of epidermal signal, using the XY projection *P*^*Π*^ of a point cloud *P* (*P*^Π^ = {(*x, y*): (*x,y,z*) ∈ *P*}) and considering only the nuclei labeled as *L*1 (Supplementary Fig. 3C-E). For instance the projected epidermal map at the time *t*_*i*_ of a sequence registered in the reference frame of the first image of the sequence will be computed as:

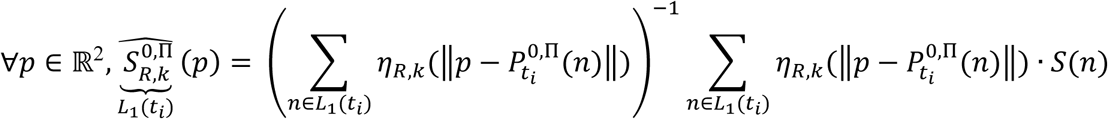

To determine the best parameter values for the function *η*_*R,k*_ we ran an extensive parameter exploration and measured the error made by mapping the signal to retain the values that yield the minimal error. This is measured by the average relative error between the actual signal value *S*(*n*) of a nucleus and the value, at the projected nuclei position *P*^0^, ^Π^ (*n*), of the 2D map computed with all nuclei but the considered one:

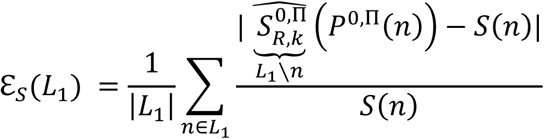

The optimal values are the ones that minimize this error over the whole available set of SAM sequences 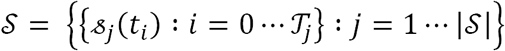 for the qDII signal, as it is our main focus in this work. The values we obtain, shown in Fig. SI IB, are *R* =* 7.5μ*m* and *k\** = 0.55μm^-1^:

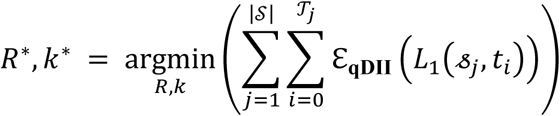

From now on, we will consider that, if not explicitly mentioned otherwise, the maps referred to are 2D projected maps computed on all the epidermal nuclei using the optimal parameters. To simplify the notations we will then write:

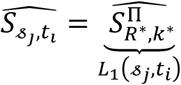

#### Sequence SAM reference frame determination

A common 3D cylindrical reference frame for the SAM sequences can be described by landmarking a set of key geometrical features for each meristem:

- the unitary vector 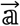 of the main vertical axis of the shoot apex
- the position 𝕔 of the apex center in the central zone (CZ) of the meristematic dome
- the unitary radial vector 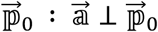 of the direction of ℙ_0_ relatively to 𝕔
- where ℙ_0_ is the position of the center of the last initiated organ primordium
- the orientation 𝕆 ∈ {–1; 1} of the phyllotactic spiral (clockwise or counter-clockwise)

We estimate the center position c on each sequence using the signal information from the *pCLV3:: C ERRY* channel quantified as 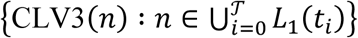. In the map 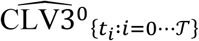 the central zone appears as a wide isotropic peak of signal intensity (Fig. SI IF). To extract the center and the radius of this peak, we threshold the map with a series of *K*_*γ*_ values depending of the average signal intensity of the sequence 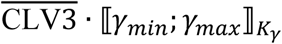 and keep for each value the largest connected component (lcc) to compute its barycenter and area:

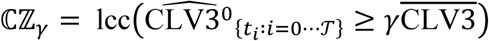

We retrieve the average barycenter and radius values of all such CZ domains obtained with *γ*_*min*_ = *γ*_*max*_ = 1.8 and *K*_*γ*_ = 7:

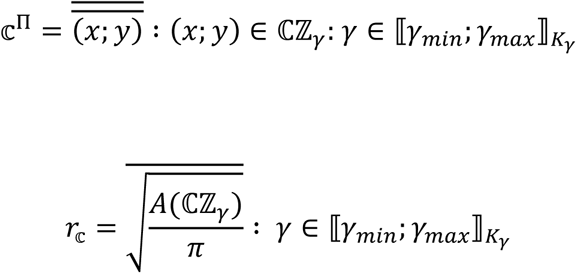

The use of a normalized CLV3 threshold made the values of *r* _𝕔_ very homogeneous among the considered individuals around an average value of 28 μm (Fig. S11G). Finally, the *z* position of the CZ center is estimated using the local *z* information:

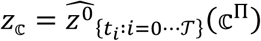

This center can be placed before estimating the main vertical axis 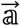 only because the tissues are imaged from above and we can make the approximation that 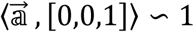. In any case we try to position of this axis by applying a small rotation *R*_*x*_(*ψ*_*x*_) *Ry*(*ψ*_*y*_) around the x and y axes to the sequence registered positions centered on 𝕔 *P*^𝕔^ *= P*^0^ *—* 𝕔. This transformation leaves 𝕔 unchanged. We note:

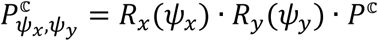

Each rotation is associated with a main axis vector 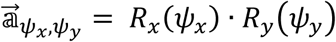·[0,0,1]. Given the radial symmetry of the meristematic dome, the optimal axis should be the one for which the function 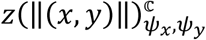 shows less variability, at least up to the radial distance where the dome ends. We take this extent into account by applying a Gaussian weight of radius *σ*_*r*_ = 20μ*m*, function of the distance *r* = ‖(*x,y*)‖ of the rotated points to the center. We look for the optimal values of the rotation angles with the simple constraint that 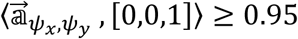 by evaluating the error to the local mean *z* of the rotated points:

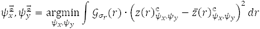

We consider that 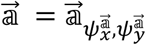 and note 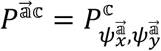. The next step is to locate the 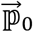 direction. This can be seen as a 2D problem if we consider the points 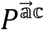, for which the *xy* plane is orthogonal to 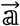. If we consider a polar coordinate system centered on 𝕔 for this plane(*r, θ*), we are ultimately looking for the angular coordinate 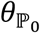 of the direction of the 𝕡_0_ primordium. In Supplementary Note 1, we detailed why 𝕡_0_ should correspond to a maximal auxin concentration in the meristem. Therefore, we look for the minimal value of the 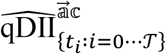 map, restraining the search to r ∈ [*ρ*_*min*_*r*_𝕔_; *ρ* _*max*_*r* _𝕔_] to avoid artifactual detections (we used *ρ*_*min*_ = 0.9 and *ρ*_*max*_ = 1.4 in the analysis)

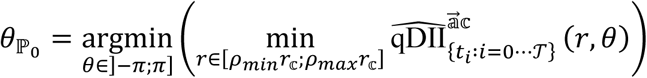

Finally, instead of estimating the orientation from the data, we chose to rely on manual expertise to determine visually the clockwise or counterclockwise orientations of the meristems. This produces a simple identity or reflection matrix, depending on the case:

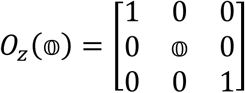

#### Aligned LI nuclei points

In the end, the determination of the SAM landmarks 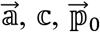 and 𝕠 allows to transform the sequence registered points *P*^0^ into a common 3D reference frame in which we will be able to compare different individuals locally. We note *P*\* the positions of L1 nuclei points in this common reference frame:

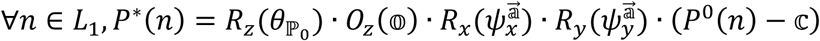

**Aligned LI image slices**

We also use the obtained transform to register the original images into the common reference frame by applying the same transform expressed in homogeneous coordinates:

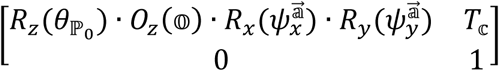

This registration is performed using the block matching computational library(*15*) and creates a registered image 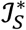 expressed over a voxel grid *I*\* that is centered on 0 in x and y and slightly shifted in z so that:

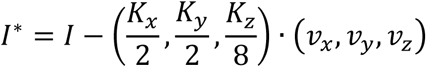

We used the 2D aligned epidermal maps in order to create the 2D projected views of these registered images displaying only a single layer of cells that can be seen in Fig. 2A-C, Fig. 3A-B, Fig. 4A and Fig. 5A-H. More specifically, we use the *z*\* coordinate of aligned *L*_1_nuclei points to compute the 2D map 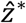 over the centered grid coordinates *I*\*, and produce a 2D image defined over the same xy grid as the original one but keeping the voxel value of one z slice per pixel:

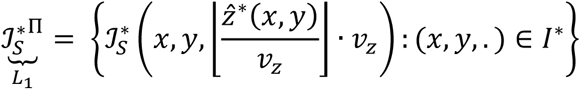

The produced 2D image displays the intensity levels of a single curved image slice that goes through the nuclei of first cell layer, making information appear more clearly than a simple maximal intensity projection that would also include intensity from the inner layers.

#### Organ primordia 2D detection

In the aligned SAM reference frame, we expect to find organs in comparable developmental stages at very close spatial locations for all individuals, under the global hypothesis of a stationary and regular development. Therefore we try to use some *a priori* knowledge on regular phyllotaxis to detect the positions of the ranked organ primordia in the meristem, namely 𝕡_0_, 𝕡_1_, 𝕡_2_ and so on.

Previous works on auxin dynamics in the meristem suggest that primordia correspond to local accumulation of auxin, which would be detectable as local minima in the 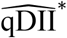 map, but also that soon after organ initiation, an auxin depletion area is formed, creating a local maximum in 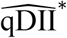. In that respect, our primordia detection procedure consists first in detecting extremal points in this map and in labeling them in a second time by organ primordium rank.

To locate extremal points in 2D, we are not only interested in absolute extremality (namely peaks and troughs of the map) but also for points that are extremal in a given direction, and we will therefore detect ridges and valleys to construct a landscape transform of the 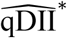 map. Peaks, which are maximal in any direction, will appear as convergence points of ridges, throughs as convergence points of valleys, and we will also retrieve saddle points as convergence points of a ridge and a valley.

Classically, extremal points are detected as zero-crossings of the gradient of a function. In our case, we discriminate between positive zero-crossings ø+ and negative zero-crossings ø^−^ that would correspond respectively to local maxima and local minima when applied on a gradient. For ID scalar function *f*: ℝ → ℝ, zero-crossings are defined as follows:

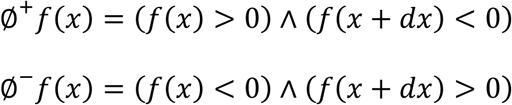

Now, if the scalar function is a 2D function *f*: ℝ^2^ → ℝ, we can define a 2D zero-crossing, checking if the function is locally changing sign along each dimension, for instance:

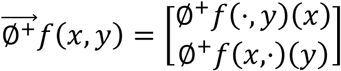

The map gradient is indeed a 2D function, but is not scalar since the gradient itself will be a 2D vector. To cope with this, we introduce the notion of (here positive) zero-crossing of a function 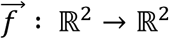?, along the direction given by the angle *ω,* written 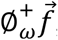, measuring if the scalar function restricting 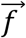 to the given direction is changing sign when we follow this direction:

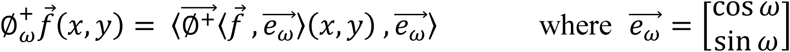

Using this local directional zero-crossing information, we can compute 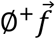 as a scalar measure of the proportion of directions in which the vector field is consistently changing sign, by integrating the zero-crossing of 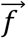 over all possible directions:

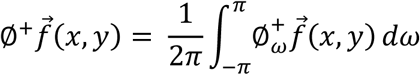

By applying this integrated zero-crossing operator, we can compute the “landscape transform” of a map (Fig. SI 1H), consisting of a local ridge estimation Λ+ and a local valley estimation Λ^-^, and obtain a way to quantify in detail local extremality on the 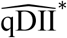 map:

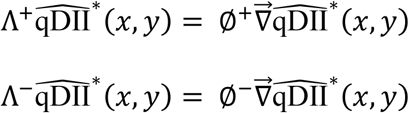

We first look for saddle points in order to eliminate them and disconnect the otherwise continuous networks of ridges and valleys. Saddle points • belong to both a valley and a ridge and are detected using their product, then valleys ▾ and ridges ▴ can be found outside the saddle locations, using different thresholds in both cases:

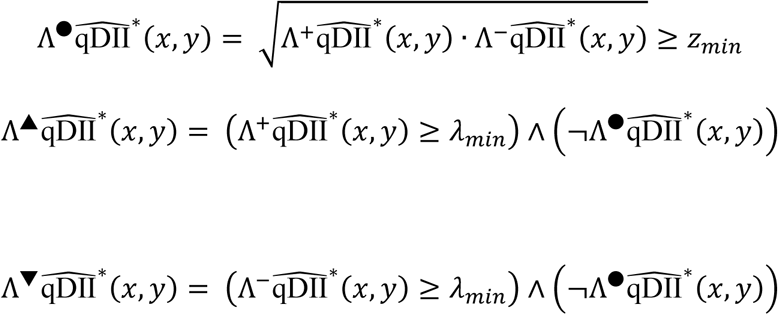

We run this binary labeling with the values *z*_*min*_ = 0.01 and *λ*_*min*_ = 0.03 and consider the connected components of each binary map as the set of extremal features of each kind, resulting in a set of regions (Fig. S11H), denoted for instance 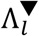 for the connected valley labeled *l.* To sum up the information, each element *I* of those three label sets written •, ▾ and ▴ is associated with a single spatial position (*r*_*l*_ *θ*_*l*_), a measure of extremality *λ*_*l*_ a value of signal 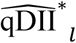 and an area *A*_*l*_ as such:

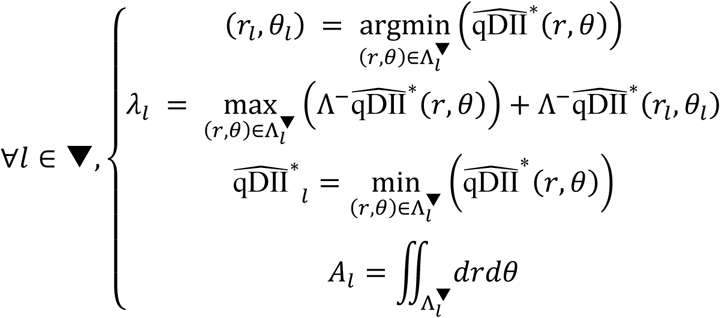

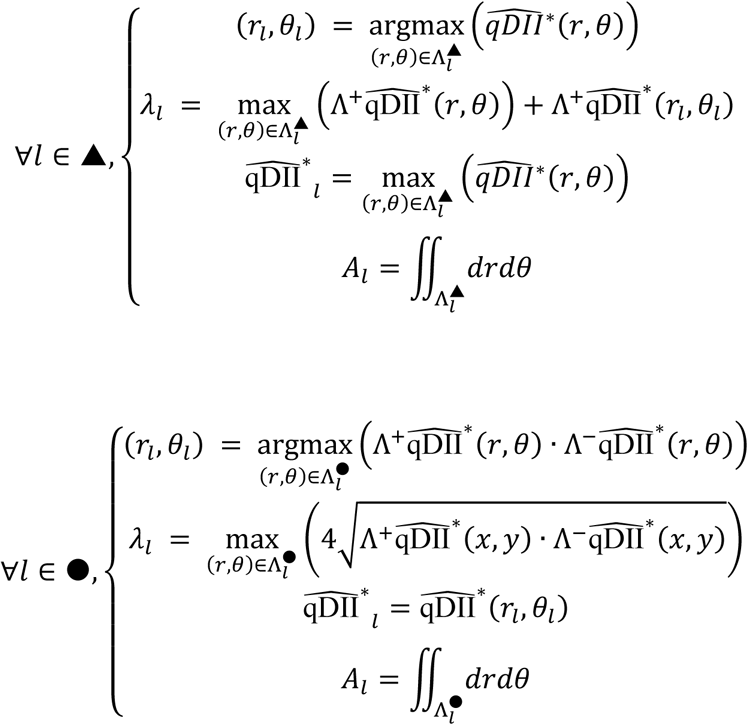

In a second time, we need to identify which of these extremal points are the ones corresponding to the youngest organ primordia. For this we use several assumptions from prior biological knowledge on the organization of the shoot apical meristem and the auxin distribution at the level of the primordium, namely:

1. Organ primordia are marked by a local maximum of auxin concentration
2. Primordia are organized in a regular spiral of divergence angle *α*\* = *2π*/*φ*^2^ ∼ 137.5°
3. In this spiral, organ primordia can be ranked by their distance to the center
4. No organ can initiate inside the central zone (CZ) of the meristem
5. The auxin depletion zone of a primordium is located between its maximum and the CZ

If we consider the organ primordium of rank *i* hereafter noted ℙ_*i*_, we look for two extremal points, a maximum of auxin (minimal point of 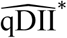) 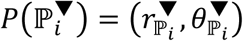 that actually defines the position of the primordium (i) and a minimum of auxin (maximal or saddle point of 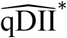) 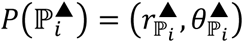 which should follow the following rules:

1. 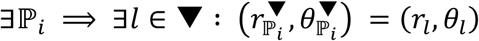
2. 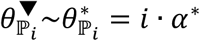
3. 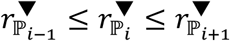
4. 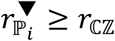
5. 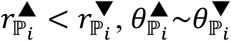

To take these assumptions into account, we perform our identification of 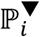 and 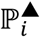 among the detected extremal points •, ▾ and ▴ by associating to each of them a positive score 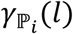 relatively to 𝕡_*i*_, measuring how good a candidate it would be for 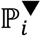 (for points in ▾) or for 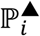 (for points in ▴ ∪ •). The idea behind the formula is to provide a result that complies as much as possible with the rules mentioned here above. The score is the product of an intrinsic, primordium independent, score *γ*(*l*) and of a spatial term 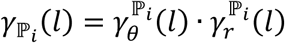 that takes into account the organization of the primordia.

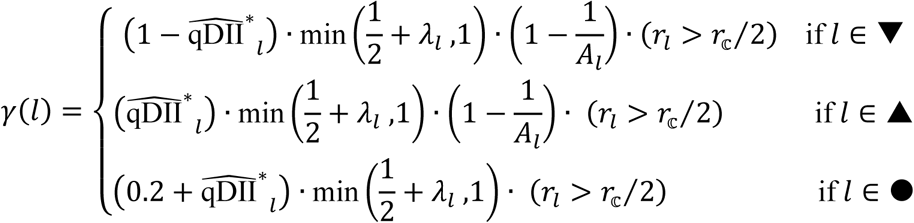

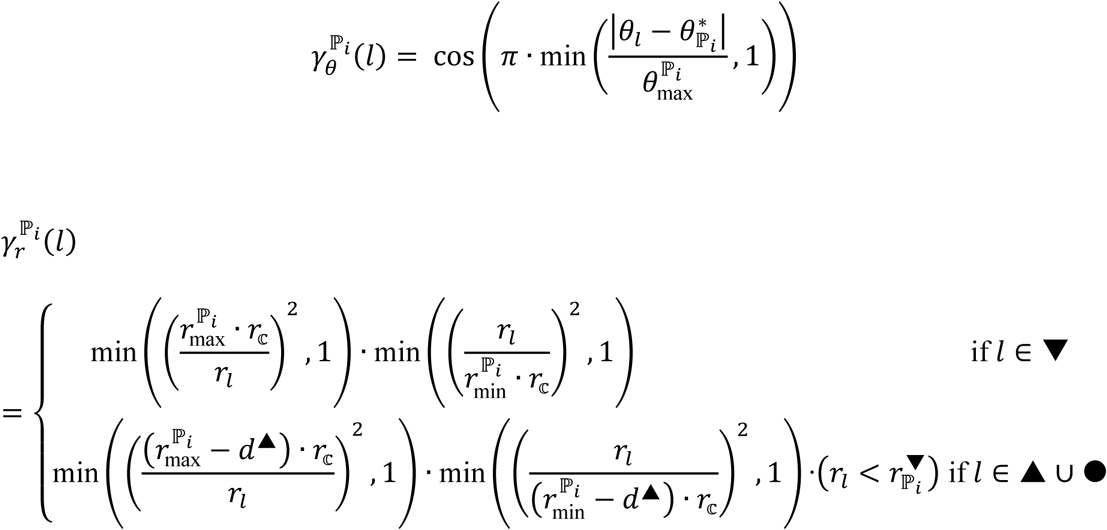

For each considered primordium ℙ_*i*_, we first identify the point corresponding to a maximal value of auxin, provided there exists one with a score value above a *γ*_min_ threshold:

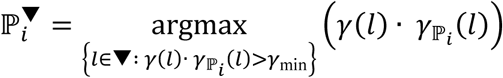

Then this detected point is used to compute the scores of minimal points of auxin by the means of 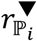 (set to +∞ of 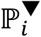 is not defined) and we identify the minimal point associated with the primordium, provided it exists:

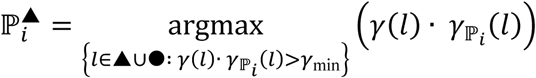

Each identified primordium extremal point is then associated with a spatial position 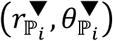 or 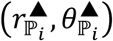 (Fig. S11I) which can be used to compute estimates of all the spatial signals (starting with 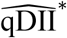) which allows us to track signals at the level of organ primordia.

On all the meristems presented in the article, we performed a manual correction of the primordium assignment of auxin extremal points, to make sure that we recover information from biologically meaningful locations. This manual labeling allowed us to perform an evaluation of the detection method, leading to an average 75.4% of correct detection (using a Jaccard index) rising to 87.3% when considering only primordia stages between 𝕡_-2_ and 𝕡_3_ (Fig. S11J).

#### Developmental Time Estimation

It is possible to position the different individual acquisitions from various time-lapse sequences onto the same developmental time axis. Making the assumption that all the considered meristems develop at a comparable rate of 2 new organs per day, corresponding to a plastochrone time of 12h, and that they can, at the first order, be considered synchronous (similarly labeled organ primordia being at comparable developmental stages) we propose a very simple developmental time indexation. For a sequence of 𝒯 acquisitions taken at {t_0_ = 0h,*t*_1,_… *t*_𝒯-1_}we choose to assign to each acquisition at *t*_*i*_ a developmental time:

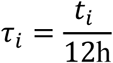

#### LI Dynamic Signal Maps

Using the indexation of individual acquisitions in developmental time and the aligned L1 signal maps, we reconstruct a large-scale picture of the typical behavior of signals in the meristem by computing averages among individuals and interpolations over time.

At the scale of one time-lapse sequence 𝓈_*j*_, for which we have computed the 2D aligned signal maps 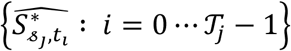 we approximate the continuous time signal by diffusing the information in developmental time by the same method we used to diffuse signal in space when computing continuous maps. At any time position *τ*, the estimated map is a weighted average of single time maps, where the weights are time-distance-based density coefficients computed using the *η*_*R,k*_ function with a time radius *R*_*τ*_ and a time slope *k*_*τ*_:

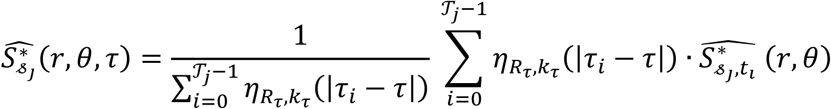

However, this approximation only enables us to reconstruct the evolution of signals over a time range equivalent to the duration of acquisitions. To extrapolate furthermore and obtain a more complete view of primordium development, we use the spatio-temporal periodicity of the system to consider the information at different spatial locations as information at different temporal positions. More precisely, we use the fact that to look *p* plastochrones further in time is equivalent to rotate the system of the angle 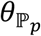 corresponding to the direction of the primordium 𝕡_*p*_. Therefore we use the angles 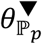 estimated with 2D primordium detection to apply rotations on the computed maps for primordia stages going from 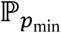 to 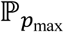 and therefore compute a dynamic map that covers a much larger temporal range (the first formula becoming a particular case with *p*_min_ = *p*_max_ = 0):

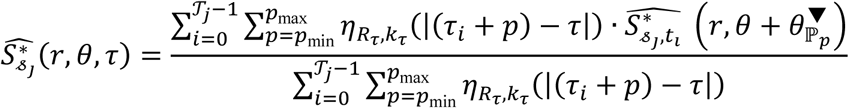

Finally, using the fact that all 2D signal maps from different acquisitions are registered into the same reference spatial frame, we reconstruct the average map over a population of meristems by generalizing this formula over the whole available set of time-lapse sequences *𝒮* = {*𝓈*_*j*_: *j* = l … |*𝒮*|} to combine all the quantitative spatio-temporal information into one dynamic map reflecting the canonical behavior of the system:

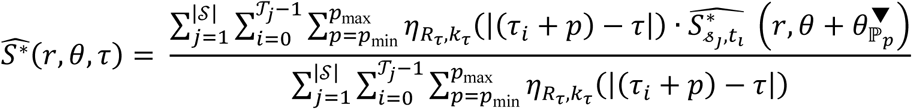

This computation results in dynamic 2D maps as those presented for Auxin signal in Supplementary Video 1. These were obtained with parameter values *R*_*τ*_ = 0.1, *k*_*τ*_ = 2, *p*_min_ = –3, *p*_max_ =5.

**Figure S11.**
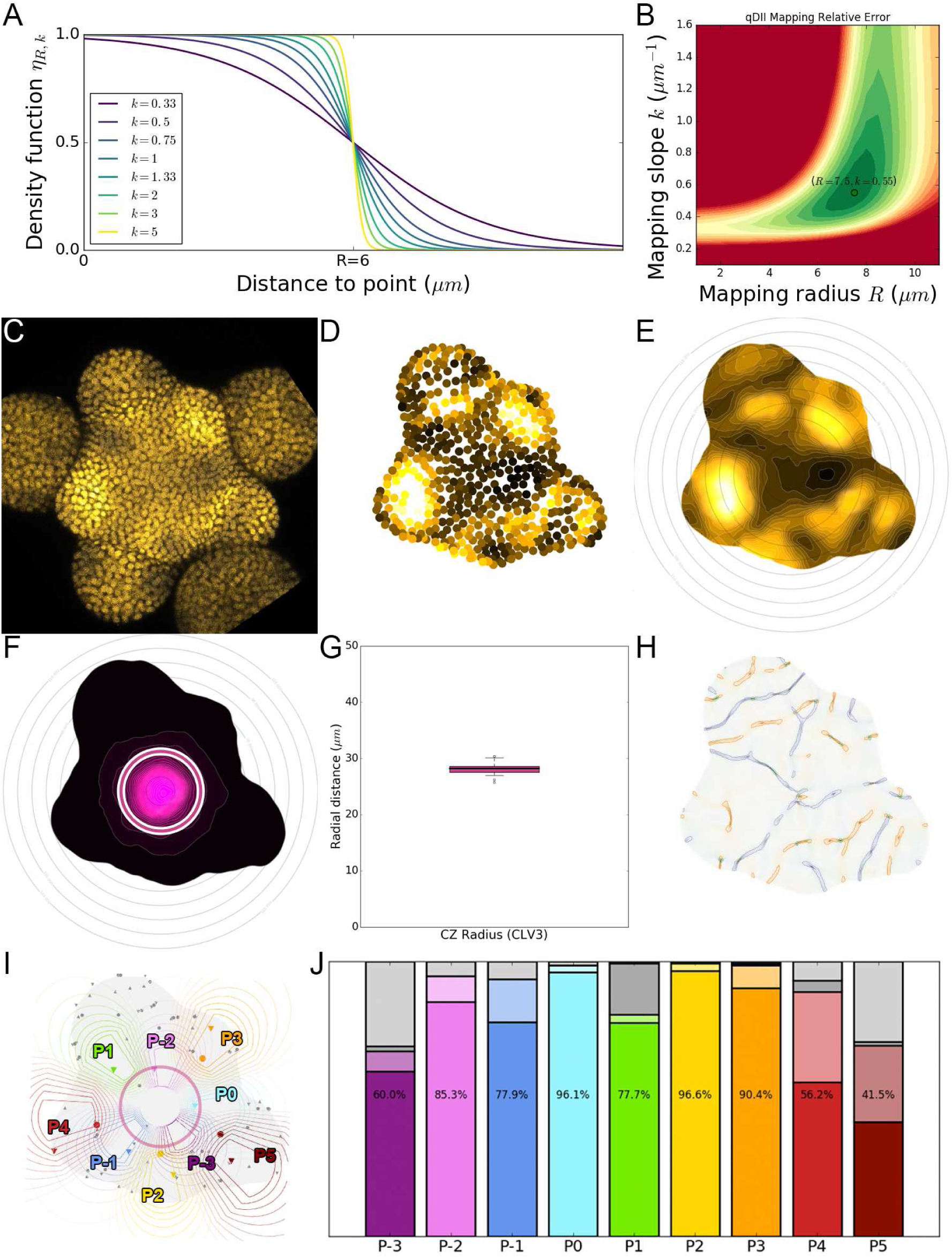
Using 2D continuous maps of epidermal signals to identify SAM landmarks. **A.** To build a continuous 2D map, we diffuse the signal in space by computing a local average of discrete signal values using a kernel function whose extent and sharpness are set by twoparameters *R* and *k.* **B.** Optimal values of these parameters were determined on the signal of main interest qDII and are the ones used throughout the analyses. **C-E.** Using the L1 nuclei detected in the confocal image **(C)** and their quantified signal values projected in 2D **(D)** we compute a 2D map of epidermal signal **(E)** in this case qDII. **F.** Using such a map computed on the quantified CLV3 levels, we are able to locate precisely the center of the meristem and estimate the extent of the CZ. **G.** The extents of CZ estimated using the CLV3 maps show a very limited variability around the 28µm value among the observed individuals (N=21 SAMs). **H.** Based on qDII maps a detection of extremal features (ridges, valleys and saddles) is performed and extremal points are extracted. I. Assuming a minimal regularity in position w/r to CZ and divergence angles close to 137.5°, the extremal points are given a score relatively to each considered primordium rank, and a maximum of one auxin maximal point and one auxin minimal point is assigned to each primordium. **J.** Comparison of automatically detected auxin extremal points with expertized ones demonstrate a very accurate detection between ranks P0 and P3, with a decreased performance when the features are less well defined (no absolute minimum before P0, several maxima after P4). Color indicates the rank of primordia. Filled color indicates accurate detection, light color indicates correct detection but inaccurate location, dark grey indicates false negatives, light grey indicates false positives.

### 2. Membrane marker image and PIN1 polarity quantification

#### Automatic segmentation of membrane images

In order to be able to quantify membrane-localized signal, we need a segmentation of the tissue at the cellular level. Such automatic cell segmentation procedures are often limited by their capacity to detect the right number of “seeds” prior to the segmentation of the image according to membrane localized signal.

In our case, the presence of a constitutive nuclei targeted signal (*pRPS5a∷Ta BFP*), allows to compute the nuclei coordinates in the image. It is thus possible to use these coordinates as seeds to initialize a watershed segmentation algorithm. The quality of the obtained segmentation, in terms of “correct number of cells detected”, is then directly linked to the nuclei detection quality. Compared to parametric seed detection by methods such as local minima detection (by h-transform algorithm) followed by connected component labeling, the use of detected nuclei signal coordinates allows to reduce the over and under segmentation problems (data not shown). To summarize the pipeline used for this automatic membrane segmentation step, we performed:

1. An adaptative histogram equalization (*18*) for all z-slices of the membrane stacks to improve and normalize the contrast;
2. Isometric resampling to a voxelsize of (0.2, 0.2, 0.2(μm), when original images are (0.2, 0.2, 0.5μm), to performs Gaussian smoothing and obtain smoother segmentation in Z;
3. Gaussian smoothing of the membrane intensity image, with *σ* = 0.2*μm* to reduce noise in the image when performing watershed segmentation;
4. Create a “seed image” from the nuclei coordinates to initialize the watershed algorithm;
5. Run the seeded-watershed algorithm with isometric smoothed intensity image and seed image.

The steps 2 to 5 correspond to the pipeline described in(*16*) where more technical details can be found. No post-segmentation corrections where performed, no cell-fusion (in case of oversegmentation) or morphological corrections (median filters to smooth the walls). In the end we obtain a segmented image *𝒮* that assigns an integer label to every voxel of *I.* The cells of the tissue are represented by independent connected regions of voxels (so that the same label can not be assigned to voxels that are not part of the same connected component of *𝒮*). The background corresponds to a specific label that is systematically set to 1 to ensure consistency between images. Each cell labeled *c* > 1 is the then represented by a connected region *𝒮*_*c*_ so that:

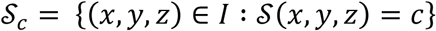

#### Cell barycenter extraction

To obtain the cells barycenter, we used the SciPy library (*19*) and the center of mass function that simply estimates the position *P*(*c*) of the center of the cell labeled *c* in the segmented image as the average of the voxel coordinates of the cell region *𝒮*_*c*_:

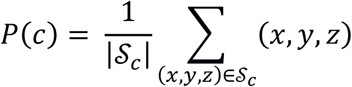

#### Cell layer estimation from segmented images

To be able to automatically determine to which layer a given cell belongs to, we use the topology of the tissue, notably the “background region” *𝒮*_1_. Indeed, the biological definition of the epidermis, also called layer-1 or L1, is to be in contact with the outside world. We therefore label as *L*_1_ any cell *c* such that its region *𝒮*_*c*_ can be considered a neighbor to *𝒮*_1_.

Neighborhood relationship is defined through the notion of surface of contact between two regions in he segmented image. In a first time, we consider the 6-connectivity to define neighborhoods at voxel-scale. However, to be robust to potential segmentation errors, we estimate the area of all surfaces of contact representing walls between two cells *c* and *c*’, and consider as neighbors only those for which the area is greater than a given threshold *A*_min_. We compute the area of contact 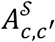 as the sum of surfel areas between pairs of 6-adjacent voxels so that one is labeled *c* and the other *c’.* An analysis of the distribution of wall areas (data not shown) led us to consider that a value of *A*_min_ = 5μm^2^ would be suitable. We note *N*(*c*) the neighbors of the cell c, namely the labels that verify the surface of contact condition:

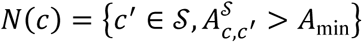

Therefore, we obtain *L*_1_ = *N*(1). Subsequently, we can define the cells belonging to the second layer (*L*_2_) as those in contact with the *L*_1_ also filtered by the same minimal wall area: 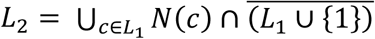.

#### Reconstruction of LI anticlinal walls

In order to quantify the PIN signal intensity at the level of each individual cell membrane, we need to have a precise identification of the cell walls and a faithful 3D representation to describe their position and orientation. If the boundary between two cells can be extracted as a set of voxels in the segmented image *𝒮,* it will generally be too sensitive to noise and to image resolution to be used as such. We chose therefore to use a triangular mesh representation with a high resolution to represent accurately the cell walls.

To obtain such meshes, we apply the Marching Cubes algorithm(72) to each cell *c* represented by its connected region *𝒮*_*c*_of identically labeled voxels in the segmented membrane image *𝒮.* This produces a triangular mesh ℳ_*c*_, generally closed (except on image borders) and with voxel-like resolution. This mesh represents the shape of the cell by a set of vertices with 3D coordinates *P*_*c*_ and a set of triangular faces linking those vertices *T*_*c*_: *ℳ*_*c*_ = 〈*T*_*c*_, *P*_*c*_〉.

Using Marching Cubes ensures us that, in an 8-voxel cube with only two labels *c* and *c’* (which typically occurs at the interface between two cells) the algorithm will create the same vertices whichever label is considered as 1 or 0. In other words, we know that two cells that are neighbors in the image (with a large enough surface of contact) will have common vertices in their mesh reconstructions ℳ_*c*_ and ℳ _*c’*_. We use this property to construct the cell wall mesh 𝒲_*c,c’*_, = 𝒲_*c’,c*_ of the interface between *c* and *c’* as the restriction of one of the two meshes to the set of common vertex points *P*_*c*_ ∩ *P*_*c’*_,:

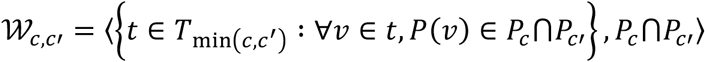

Each wall mesh undergoes then a phase of triangle decimation(*13*) and isotropic remeshing(*14*) to obtain a regular surface so that the typical length of a triangle edge is close to 0.5μm, which is about the voxel characteristic dimension. On the triangular mesh, we estimate the normal vectors 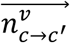 at each vertex, ensuring they all point from *c* to *d,* and the area of each triangle that allows us to estimate the total area *A*_*c,c*’_ of the interface between *c* and *c’.* Note that the Marching Cube intersection, along with the decimation and smoothing, will produce a mesh that is smaller that the actual wall (the intersecting part does not extend to cell edges) and the wall area will be underestimated. On the other hand, voxel-based area estimation is known to be largely overestimating, which ultimately provides a way to have both lower and upper bound estimates of the wall areas.

#### Quantification of PIN signal at wall-level

We consider that the PIN polarity ℙ_*c→c′*_ = ℙ_*c′→c*_ of a given cell interface, i.e. whether the PIN efflux carriers orient the flow of auxin from *c* to *c’* or from *c’* to c, is given by the differential of PIN concentration that exists between the plasma membranes of *c* and *c’* at their interface. Our way to access this information is through the difference of PIN signal intensity in the image on either side of the cell wall marked by the PI signal intensity around 𝒲_*c,c′*_. We designed a method that relies on the sampling of both PIN and PI image signals across the wall to make a decision on whether there exists a significant difference in PIN intensity levels, and therefore a polarity of the wall.

The vertices and normals of the triangular mesh 𝒲_*c,c*’_, are used to generate a set of 3D cylinders in which we will sample the image signals (Fig. S12A). We chose to exclude the vertices located on the contour of the mesh as the normal estimation may be less robust when the vertex is not fully surrounded by faces, and only consider the inside of the mesh 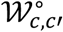, At each inner vertex *v,* the main axis of the corresponding cylinder 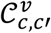 is given by the local normal vector to the mesh 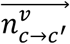 and is characterized by its radius *r*_*𝒞*_ and its extent on each side of the wall *d*_*𝒞*_ (Fig. S12B-D). In the following we will consider only the image voxels lying inside this cylinder, which, if we note(x,y,z)^Π^ the projection of a voxel (*x,y,z*) on the main axis 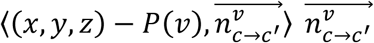, can be defined as:

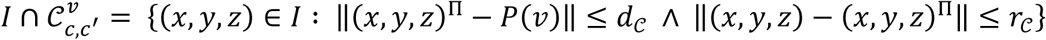

We project the signal values of these voxels on an 1-dimensional axis assigning them a signed abscissa 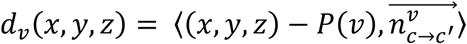, which allows us to consider the spatial distribution of signal intensities along the cylinder axis (Fig. S12C). On this axis, the position *d*_*v*_ = 0 corresponds to the intersection of the cylinder 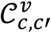 with the wall mesh 𝒲_*c,c*’_. This position might be shifted from the actual cell wall due to the stacking of residual errors from the consecutive processing steps (sequence segmentation artifacts, meshing simplifications, smoothing approximations). The idea is therefore to come back to the actual membrane-marker image signal 𝒥_*PI*_(*x,y,z*) to locate the position of the cell wall in the cylinder with a higher precision.

We achieve this precise location by estimating the abscissa of the mode of membrane signal intensity from the distribution 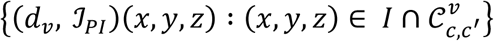. This is done by fitting a Gaussian-shaped function, optimizing its parameters to minimize the squared error to the distribution (Fig. S12E). The function uses 4 parameters and its form is given by:

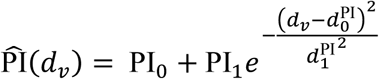

The least-squares estimation of the function notably gives us a value 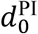 that marks the estimated abscissa of the PI maximum within the considered cylinder, which we will use as a reference defining the actual position of the cell wall. Then, the signals are quantified on either side of this abscissa, up to a fixed distance *d*_max_, by computing the average voxel intensity (Fig. S12F). For instance, we determine the two values of PIN signal intensity as follows:

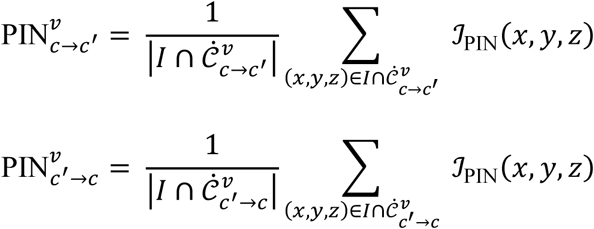

where we note 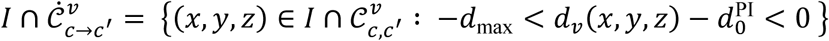 and 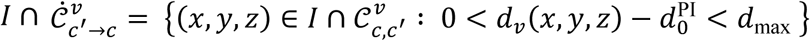. Note that the definition is symmetrical, meaning that the result would be exactly the same if *c* and *c′* were to be permutated.

**Figure S12.**
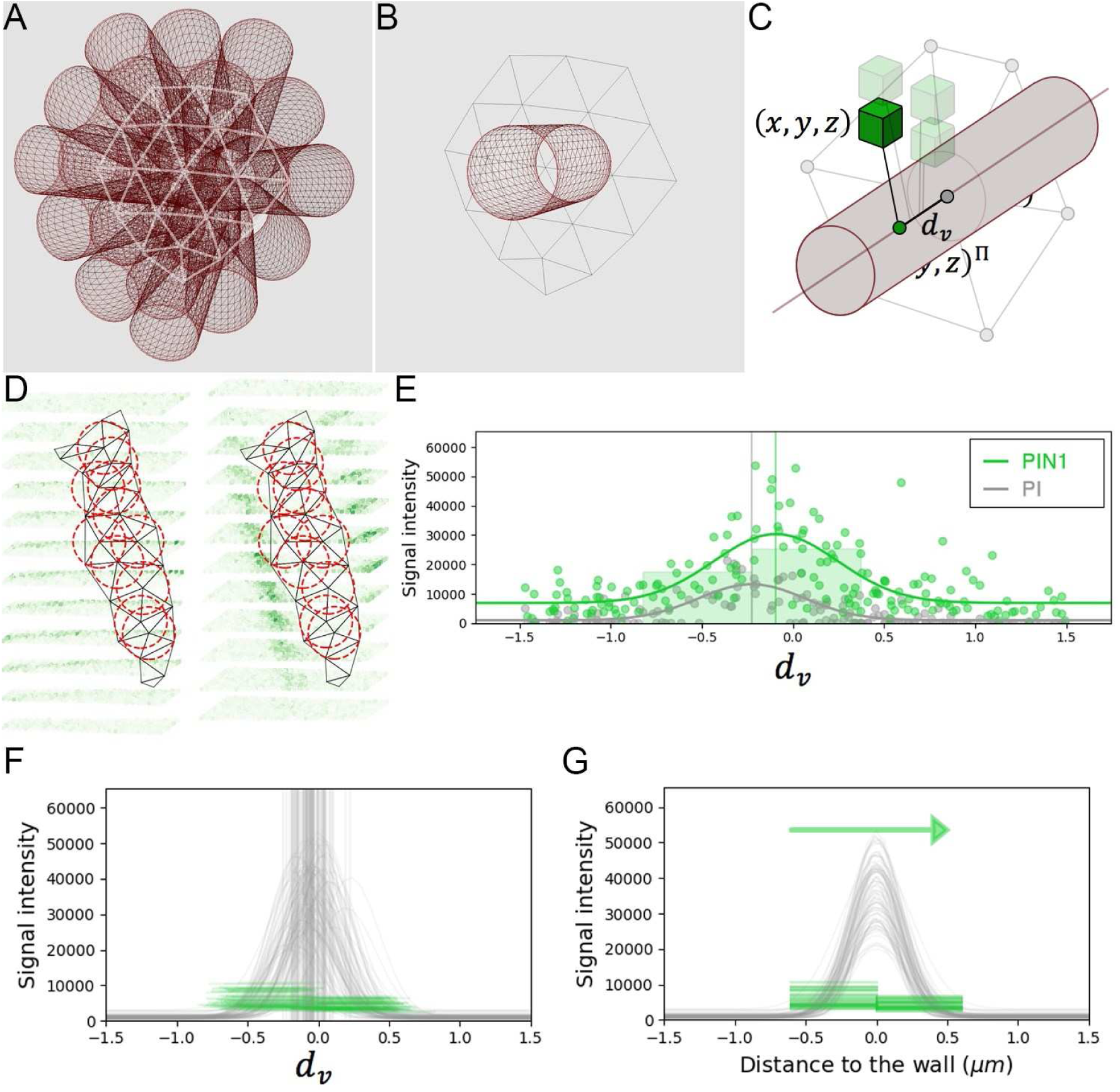
Cell wall-level estimation of PIN polarity. **A.** The triangular mesh representing the wall is used to generate a set of 3D cylinders locally orthogonal to the wall and placed at each vertex of the inside of the wall. **B.** The radius of the cylinders is close to the typical resolution of the mesh. **C.** For each cylinder, all the neighboring image voxels are projected onto its main axis, and kept if within the distance defined by the cylinder radius. Distance of the projected voxel to the wall vertex is used as abscissa for the evaluation of 1d polarity **D.** The set of all wall cylinders allow sampling the PIN image signal on either side of the cell wall at different locations. **E.** The **1**-dimensional distributions of both PI and PIN image signals along each cylinder are used to locate precisely the cell wall position and quantify signal levels on either side. **F.** PIN levels are estimated left and right of the detected wall abscissa up to a fixed distance of 0.6(μm. G. Significant difference between left and right distributions of PIN levels across the wall allows deciding for local PIN polarity at the scale of the cell wall.

By performing this two-sided estimation on every cylinder defined by the wall triangular mesh, we end up with two parallel signal distributions 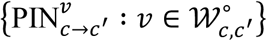 and 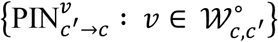. We test statistically whether these distributions can be seen as significantly different by an ANOVA test, and decide that a polarity exists when the test gives a p-value < 0.05 (Fig. S12G). In such case the polarity is given by the sign of the difference between the medians PIN_*c*→*c′*_ and PIN_*c*→*c′*_ of the two distributions:

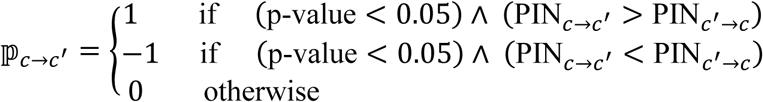

Note that, once again, the definition is symmetrical ensuring that ℙ_*c*→*c′*_ = ℙ_*c′*→*c*_. In any case, we also define the wall-level intensity of PIN signal as the average of the signal medians of either side:

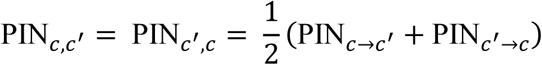

Finally, each cell wall is also characterized by its barycenter *P*_*c,c′*_ and by a single normal vector computed as the average of the inner wall normals:

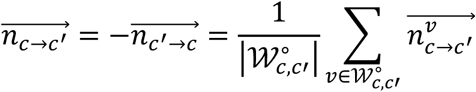

#### Computation of cell polarity vectors

The local wall polarity needs to be integrated to the cell level if we want to describe the local directionality of PIN carriers inside the tissue. To do this, we define the polarity of a cell *c* as a 3D vector 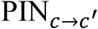 that takes into account all the anticlinal cell walls surrounding the considered cell. Each wall contributes to the resulting vector proportionally to its area, and if polarized, adds up a directional flux parallel to its normal vector with an intensity equal to the difference of PIN intensities on its sides:

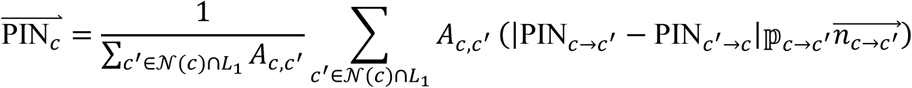

#### Alignment and map generation

To make it possible for the data computed on membrane images to be mapped onto the data from the nuclei channels of the same acquisition, we need to transform the quantified information into the same reference frame. The processes of rigid time registration and sequence SAM reference frame determination ultimately define a single rigid transformation matrix *T\** per acquired time point that transforms the physical image coordinates into coordinates in the common reference SAM frame. Therefore we simply need to apply this transform to the wall centers 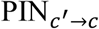, to the cell barycenters ∀*c* ∈ *L*_1_ *P\**(*c*) = *T* P*(*c*), and to apply the rotation component *R\** of this matrix to the cell polarity vectors 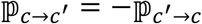.

Using the epidermal map formalism, introduced in 2D maps of epidermal signal, and from these transformed coordinates, we can then compute an aligned continuous map of PIN intensity based of the wall-level quantification of wall signal and the estimation of wall areas:

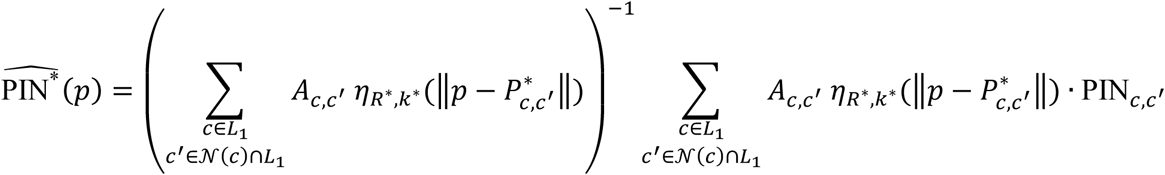

We also use the same method to compute an aligned vectorial map of PIN polarities, this time based on the estimation of cell polarity vectors:

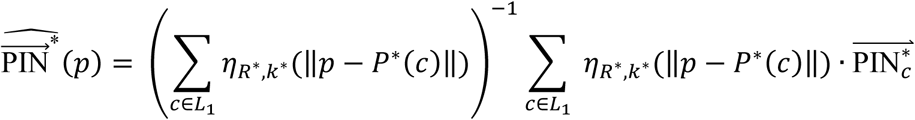

In particular, this is the data we use to quantify the convergence of PIN directions at tissue scale by applying the divergence operator to the vector field resulting from the map computation 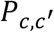. This operation gives us a scalar map where negative values correspond to areas of local convergence of the PIN directions:

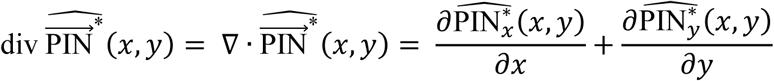

Practically, this scalar map is computed from the estimation of the vector field on a regular 2D grid of coordinates by applying a 1-D Gaussian derivative filter of standard deviation σ_div_ = *R*/*2 to each component of the vectorial map in the adequate dimension. We use this computed map to approximate the values of div 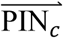 at punctual locations such as nuclei points *P*(*n*) or primordia extremal points 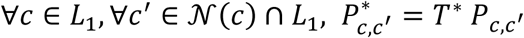 or 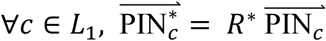 by averaging the values at the 4 closest grid coordinates. Indeed, it would be very complex to estimate this quantity on a discrete set of points from the 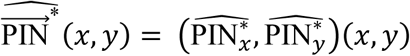 vectors, on which the divergence operator would be non-trivial.

#### Supplementary Method 3. Extrapolated cell motion in the developmental continuum

Throughout this work, we strongly relied on the spatio-temporal periodicity of phyllotactic systems that we demonstrated to be a valid assumption for our considered SAMs. Indeed, the high angular precision and limited plastochrone variability make the observed systems close to a steady developing regular phyllotactic system with a divergence angle *α =* 137.5 ± 6.7° and a plastochrone *T =* 12 ± 2h (Fig. S2D-G, Supplementary Method 1).

Notably, we considered that, in the 2D cylindrical reference frame centered on the CZ of the shoot apical meristem, the dynamics of any quantifiable signal *S* must follow the properties of a spatio-temporally periodic function of spatial period [0, –*α*] and temporal period *T:*

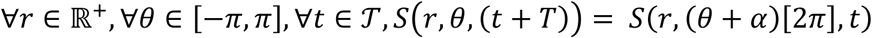

This helped us consider signal dynamics on durations that largely overpass the observation range of 10 to 14 hours, by applying successive rotations of the meristems to simulate the passing of time. Notably, an aligned SAM observed at *t* =0h rotated of 137.5° clockwise has been shown to be the best next frame to the same aligned SAM observed at *t* =10h (Fig. 1G, Fig. S2H). More generally, any primordium of stage *p* visible at time *t* can be use to infer information at time *t* + *pT.* This means that we were able to reconstruct trajectories of signals at a given position (*r, θ*) over a duration of up to 9 plastochrones (∼100 hours) from observations spanning only one, but with 9 visible primordia (P_-3_ to P_5_). This was achieved only by interpolating rotated aligned information (Movie S1).

Unfortunately, this global reconstruction heuristic could work only while we were looking at the same location in space, where the spatio-temporal property holds. To some extent, it can be generalized to robust primordia landmarks, such as auxin maxima, that we assume to be unique while moving in the course of primordium development. If they can be identified, and associated with a primordium of stage *p*, then they can be positioned on the same developmental axis at a time *t + pT* to reconstruct a developmental history at the level of this time-tracked landmark.

However, the moment we are interested in cellular processes, such as the dynamics of transcriptional response to auxin for instance, the reconstructed long-term trajectories cannot be used to draw relevant conclusions, as they reflect the dynamics either at fixed coordinates or at non-cell-specific landmark points. It is therefore necessary to find a way to access temporal cell-level information. Individual cells can be tracked in time-lapse sequences, either manually or automatically, which could be use to obtain signal trajectories over 10 to 14 hours (Fig. 2G). But to achieve the mentioned 100-hour reconstruction, spatio-temporal periodicity has to be used at some point.

#### Extrapolated tissue area tracking

We assume the cells in the central and peripheral zones (CZ and PZ respectively) of the SAM to have essentially an outward radial motion that accelerates as cells exit the central zone. This has been confirmed by the cellular motion vectors estimated from vector fields of image deformation (Supplementary Method 2) in which the azimuthal component is in average close to 0, with a limited amplitude compared to the radial component (Fig. 13SA). We note 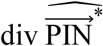 the speed of the nucleus *n* in the 2D polar SAM reference frame:

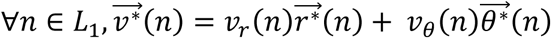

where 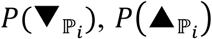 and 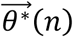 are the local normal and tangential unitary vectors at *P\**(*n*). In the following, we will then assume a pure radial motion of cells in the L1 of the SAM, i.e. that:

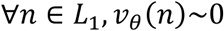

We compute the local radial cellular speed as a 2D continuous map on the L1 (Fig. S13B-C), using the same parameters as for the other cellular signals:

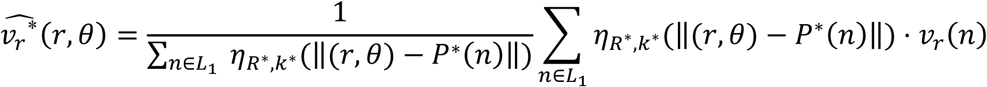

Defined this way, local radial cellular speed is a tissue-level information, that is not attached to a cell but to a spatial position (*r,θ*). Therefore it has a spatio-temporal periodicity property and we can write:

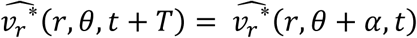

We use this spatio-temporal periodicity property of local cellular speed to extrapolate cell motion over time from a series of acquisitions of SAMs at discrete times {*t*_0_ < ⃛ < *t*_*n*_ *< T*}. Let us consider a cell with an initial position *P*(*t*_0_) = (*r*(*t*_0_), *θ*_0_) setting *r*(*t*_0_) = *r*_0_. Using acquisitions at *t*_*0*_, it is possible to estimate 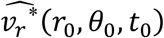, which we use to estimate P(*t*_1_) = (*r*(*t*_1_), *θ*_0_) assuming a linear motion between *t*_0_ and *t*_1_:

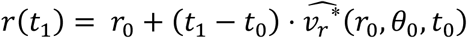

More generally, with observations at *t*_*i*–1;_ we derive *P*(*t*_*i*_) with:

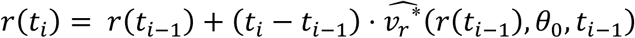

We perform this progression until we compute the last position *P*(*t*_*n*_) for which motion can not be estimated (as there is no next image it compute image deformation) However, we can still extrapolate the lastly computed motion to reach one plastochrone, and estimate *P*(*t*_0_ *+ T*) with:

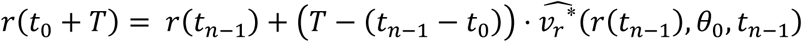

To proceed further, we would like to progress in time and estimate the cell position at *t*_1_ + *T.* This is where we use the spatio-temporal periodicity property to derive that:

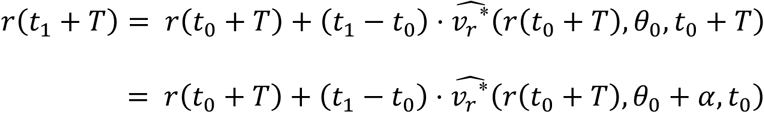

We are therefore able to estimate this next position from observations at *t*_0_ simply by rotating the radial speed map (Fig. S13D). Then iteratively it is possible to go further in time to *t*_*n*_ + *T,* extrapolate motion to *t*_0_ + *2T,* apply again spatio-temporal periodicity to reach *t*_1_ + *2T* and so on. Finally we obtain the two following general equations:

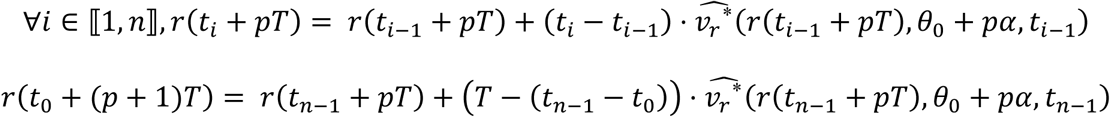

These are valid for positive integer values of *p* until reaching the maximal extent of *P* in the considered data (i.e. when the map used to estimate local radial speed becomes ill-defined), which determines a value *p*_max_. This defines a radial trajectory in the 2D space that reflects the local motion of cells along several plastochrones (Fig. S13E). Note that a rigorously identical approach can be used to go backwards in time with negative values of *p* until *t*_0_ + *p*_min_*T*, using motion vectors computed using inverse image deformation (Supplementary Method 2).

In the end, from an initial position (*r*_0_,*θ*_0_,) we obtain a discrete radial trajectory:

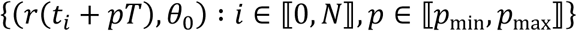

To monitor a cellular process over long time courses, the objective would be to estimate the value of the signal *S* along this spatio-temporal trajectory, i.e:

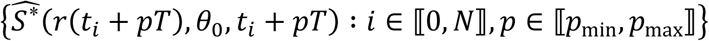

Using the spatio-temporal periodicity property of *S,* this translates into:

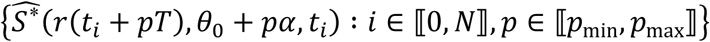

In other terms, we have defined a series of spatial locations in a time-series of acquisitions such that the sequence of signal values at these locations on the same meristem estimates the cell-level trajectory of the considered signal. If we represent them on a time-series of meristems, we define tissue areas that can be tracked, first in time then in space, to reconstruct the average behavior of a group of cells over time (Fig. S13F-H). This is the approach we use to reconstruct cell-level auxin trajectories (Fig. 2H) and to study the relationship between auxin input and transcriptional response in a consistent group of cells (Fig. 4D, Fig. 4F-H).

**Figure S13.**
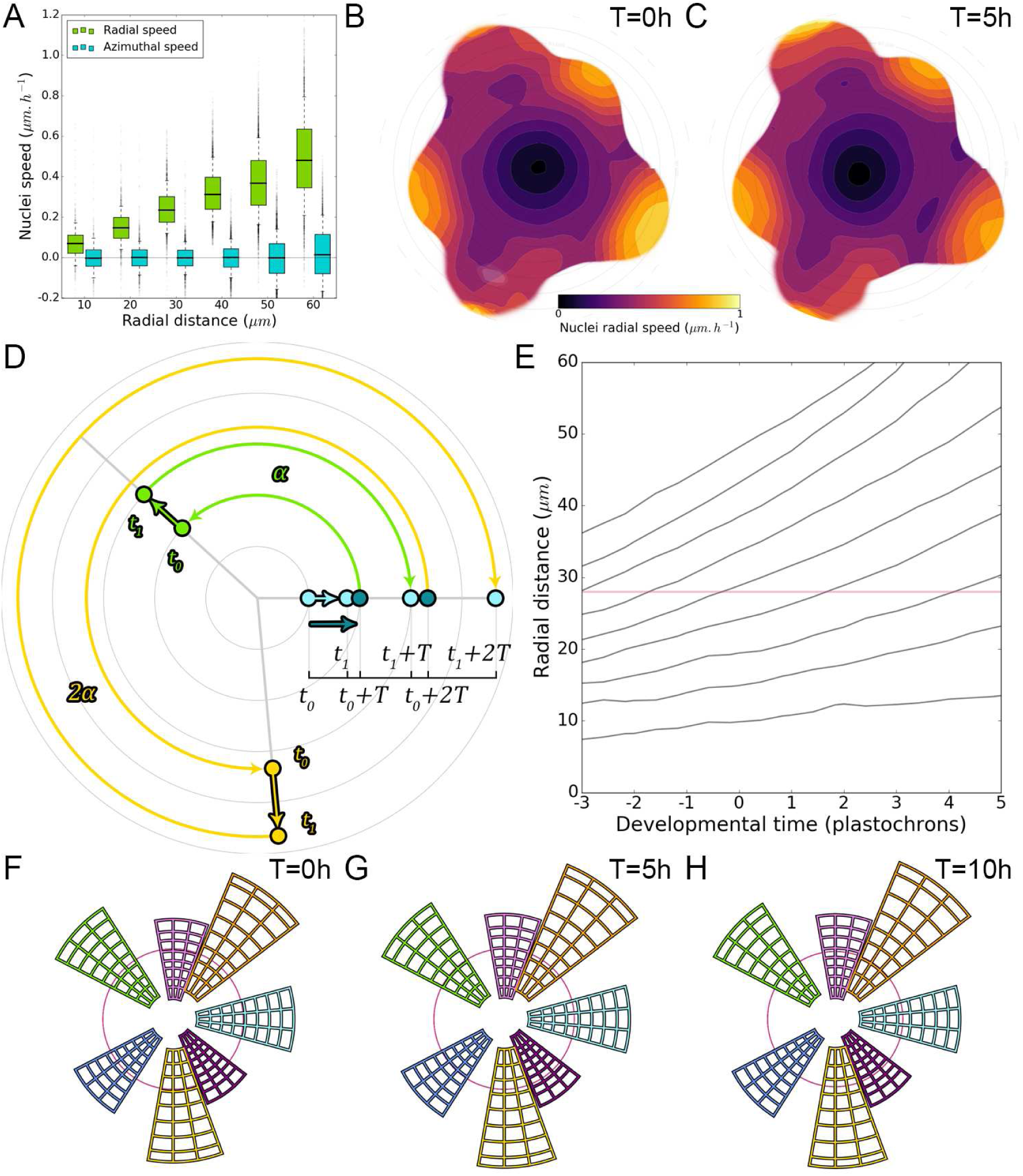
Extrapolating tissue motion to reconstruct cell-level dynamics. **A.** Cellular motion on the L1 is essentially a radial motion towards the periphery of the SAM that accelerates as cells exit the CZ. **B-C.** Average 2D maps of L1 local radial cellular speed, computed from image deformations between observations at t=0h and t=5h (**B**) and t=5h and t=10h (**C**) (N=21 SAMs). **D.** Spatio-temporal periodicity allows estimating cellular motion on several plastochrones by successive rotations: motion at to can be used to estimate position at t_1_, which can be extrapolated to t_0_+T. By periodicity, radial motion at t_0_+T is equal to radial motion at to rotated by one divergence angle α, which gives the position at t_1_+T, and so on. **E.** The iteration of this process in time allows the reconstruction of long-term radial cellular trajectories that correspond to the average motion of cells over the population over nearly 100h. **F-H.** These trajectories are used to define spatial domains that reflect cellular motion over time and allow the study of cell-level processes over long durations.

